# Dopamine in the ventral and tail of striatum supports global and local evaluation in reward-threat conflict

**DOI:** 10.64898/2026.05.01.722240

**Authors:** Isobel Green, Eshaan S. Iyer, Audrey Kang, Naoshige Uchida, Mitsuko Watabe-Uchida

## Abstract

Survival requires balancing reward seeking and threat avoidance, yet how distinct dopamine systems coordinate to support this remains unclear. Using a naturalistic foraging paradigm in which mice pursue water reward under threat from a monster object, we examined roles of dopamine projections to the ventral striatum (VS) and tail of the striatum (TS). Ablation of VS- projecting dopamine neurons impaired both distal reward pursuit and threat avoidance, with the impairment in threat avoidance paralleling effects of TS dopamine ablation. However, simultaneous recordings revealed different activity rules: VS dopamine tracked radial velocity towards the current goal as animals changed goals (reward or shelter), consistent with a temporal-difference error of spatial value, while TS dopamine encoded proximity and orientation to the threat, reflecting immediate sensory experience. Taken together, VS and TS dopamine evaluates distinct state information for avoidance. VS dopamine facilitates allocentric, goal-directed navigation, while TS dopamine facilitates egocentric, stimulus-driven threat responses.

## Introduction

Animals must maintain a delicate behavioral balance as they forage in the natural world. To avoid harm, they must prioritize defensive responses to threats, while pursuing rewards such as food, water, and warmth which are necessary to survive. Dopamine is known to facilitate learning about and pursuing rewards^1–5^. Canonical dopamine neurons in the lateral ventral tegmental area (VTA), and particularly those projecting to the ventral striatum (VS), are excited by reward and inhibited by punishment^6^. These dopamine neurons integrate value of both appetitive and aversive valences when a single cue predicts both reward and punishment^7^, consistent with the idea of a common currency for valenced stimuli, which is useful in training brain networks for value-based decision-making.

Challenging the idea of dopamine as a common currency for values, multiple studies have found subpopulations of dopamine neurons which are activated by non-rewarding or even aversive stimuli. In particular, dopamine neurons projecting to the tail of the striatum (TS) are excited by threatening and/or novel salient stimuli and facilitate threat avoidance^8–14^. From such findings in diverse settings, it has been proposed that dopamine in TS signals a prediction error of stimulus salience. This salience prediction error may be used to learn threat predictions^8–10^ and more broadly promote attention to salient stimuli, facilitating avoidance of threats (highly salient) and orienting towards surprising or potentially rewarding stimuli (moderately salient)^15^.

While TS dopamine is involved in learning from and responding to threats^13^, VS dopamine is also known to contribute to threat avoidance^16–18^, not only reward approach, raising a potential redundancy in the function of these circuits. VS dopamine neurons are inhibited by threatening stimuli and facilitate learning of negative value of threat^8,19,20^. Further, in a subarea of VS, dopamine is released by safety (e.g. when animals successfully avoid electric foot-shock), and facilitates learning of active avoidance^19,21,22^. Theoretically, therefore, animals could use negative prediction errors occurring at threat encounters and positive prediction errors occurring at safety, both signaled by VS dopamine, to learn to avoid^23–25^. Evidence of a role for TS dopamine in threat avoidance raises the question of why a separate evaluation system is needed if a value-based system already broadcasts the complete information needed for avoidance.

In this study, we examined how dopamine neurons projecting to two distinct striatal subareas influence behavior in a naturalistic foraging setting. Specifically, we tested how dopamine-mediated evaluation systems (i.e. TS and VS), both of which have been shown to be important for threat avoidance, modulate foraging behaviors under threat. One potential model is of an oppositional relationship: under threat-reward conflicts, dopamine in VS may mainly facilitate reward approach and dopamine in TS may facilitate threat avoidance, with animals using these orthogonal information streams to flexibly solve conflicts. Alternatively, dopamine in VS and TS may function in a complementary manner: dopamine in both VS and TS might contribute to threat avoidance, given the criticality of this behavior.

We found that roles for VS and TS dopamine are not oppositional in reward-threat conflicts. Animals with ablation of VS-projecting dopamine neurons decreased threat avoidance, causing better reward acquisition under threat, similar to animals with ablation of TS-projecting dopamine neurons^13^. However, their roles are not redundant, either. Activity of dopamine axons in VS was consistent with temporal difference (TD) error of spatial value, flexibly updated across contexts, whereas activity of dopamine axons in TS was consistent with threat prediction mapped on an object but not on space. Our study indicates synergistic functioning of two separate evaluation systems to promote adaptive behavior under approach-avoidance conflict with TS dopamine facilitating immediate responses to threatening objects, and VS dopamine facilitating sustained responses to both reward (approach) and threat (avoidance). These results indicate that VS and TS differ not only in their dopamine responses to valenced stimuli, but also in the state information they evaluate and the action programs they promote. This synergistic functioning appears integral to adaptive functioning under reward-threat conflict, as loss of either channel disrupts the balance between approach and avoidance behavior.

## Results

### Mice with ablation of VS-projecting dopamine neurons show deficits in reaching distal reward but not proximal reward

To understand the roles of dopamine in VS and TS in foraging under threat, we employed the monster paradigm, in which thirsty mice were free to forage for water reward (Figure 1); in some sessions, a “monster” threat was present (see Methods)^13^. Prior work found that ablating TS-projecting dopamine neurons decreased threat avoidance and increased reward acquisition in the monster task^13^. Here we ablated VS-projecting dopamine neurons to compare their functions in this paradigm. To ablate dopamine axons widely in VS, we injected the selective neurotoxin 6- hydroxydopamine (6-OHDA) into VS bilaterally (Figure 1B), injecting vehicle only in control mice.

**Figure 1.**
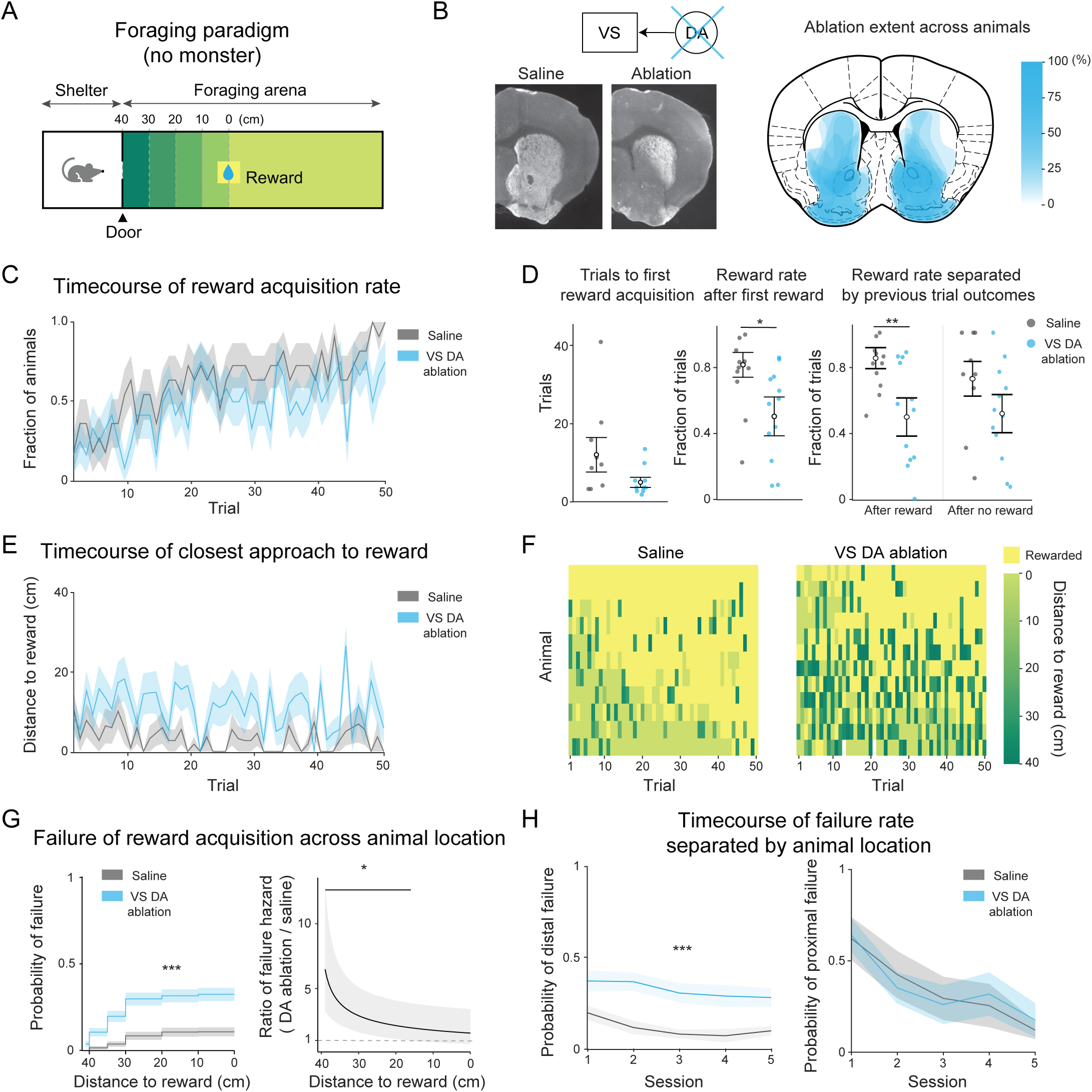
Ablation of VS-projecting dopamine neurons induces deficits in attaining distal but not proximal rewards. (A) Schematic of the foraging task. In each trial, the shelter door opened to allow a mouse to enter the foraging arena, where water is available 40 cm from the door. (B) Left, representative coronal brain sections of animals with ablation of VS-projecting dopamine neurons and of control (saline) animals. White, dopamine axons visualized with tyrosine hydroxylase (TH) expression. Right, a map overlaying the extent of dopamine axon ablation across animals. (C) Reward acquisition improved over trials, but the trajectory of improvement differed between VS dopamine ablation (blue) and control animals (gray) (Binomial GLMM: ablation: β = 0.17 ± 0.81, *z* = 0.21, *p* = 0.83; trial: β = 0.047 ± 0.010, *z* = 4.6, *p* =3.5×10^-6^; ablation × trial: β = 0.037 ± 0.012, *z* = 3.1, *p* = 0.0017). (D) Trials to first successful reward acquisition did not differ significantly between groups (left; Welch two-sample t-test: *t*(9.36) = −1.5, *p* = 0.15). In contrast, the rate of successful reward acquisition after the first reward trial was significantly reduced by VS dopamine ablation (Wald z test: middle; OR= 0.22, *z* = −2.3, *p* = 0.018). The reward rate of ablation animals was lower even after the same reward experience in the previous trials (Wald z test: after reward, OR = 0.17, *z* = −2.8, *p* = 0.0046; after no reward, OR = 0.40, *z* = −1.4, *p* = 0.15). Error bars, mean ± SEM. Dots, individual animals. (E) The distance to reward at closest approach on each trial was shorter for control (gray) compared to VS dopamine ablation animals (blue) (LMM ANOVA: ablation: *F*(1, 48.66) = 10.7, *p* = 0.0019; trial: *F*(1, 1110.59) = 7.2, *p* = 0.0070; ablation × trial *F*(1, 1114.84) = 0.031, *p* = 0.85) (F) The distance to reward at closest approach in each trial for each animal (left: control; right: VS dopamine ablation). The distance 0 (yellow) denotes rewarded trials. (G) Probability of failure of reward acquisition as a function of animal’s distance to reward differed significantly by animal group (Kaplan–Meier analysis, log-rank test: *χ²*(1) = 82.4, *p* =1.1×10^-19^). (I) The ratio of failure of reward acquisition across animal groups decreased with distance to reward (Cox proportional hazards model; Cox model ablation term: *χ²* = 26.0, *p* = 3.3×10⁻⁷; distance-varying ablation term: *χ²* = 8.3, *p* = 0.0039). (H) Failure of reward acquisition was separated by closest approach to reward into distal failure (>15cm, left) and proximal failure (<15cm, right). Ablation animals had a higher probability of distal failure (Wald χ² ANOVA: ablation: *χ²*(1) = 4.4, *p* = 0.035; session: *χ²*(4) = 4.7, *p* = 0.31; ablation × session: *χ²*(4) = 3.9, *p* = 0.41). Once animals became proximal to the reward, success rate did not differ between animal groups and improved similarly across sessions (Wald χ² ANOVA: ablation: *χ²*(1) = 0.02, *p* = 0.87; session: *χ²*(4) = 60.9, *p* = 1.8×10^-12^, ablation × session: *χ²*(4) = 3.9, *p* = 0.40). Mean ± SEM across animals. n = 11 animals (ablation), and 12 animals (control). * *p* < 0.05, ** *p* < 0.01, *** *p* < 0.001.

We first investigated the impact of the ablations on behavior in the training phase of the monster task, during which mice learn to harvest reward in the absence of the monster (Figure 1A). On each trial, the door of the shelter opened, and mice were free to enter a foraging arena to acquire a small drop of water reward. Each trial ended once the mice returned to the shelter. Animals in both the control and ablation groups steadily improved at harvesting water across training sessions, but VS dopamine ablation mice were slightly slower to improve (Figure 1C). This impaired learning rate was not due to deficient initial exploration of the arena in ablation animals, as they took similar (even slightly fewer) trials than control animals to harvest their first reward (Figure 1D left). However, ablation animals were less adept at exploiting information about this known reward: success rate after an animal’s first reward was significantly lower in ablation animals compared to controls (Figure 1D center), and the success rate in ablation animals remained lower than controls even in trials immediately following successful reward acquisitions (Figure 1D right).

We found that the reward-seeking deficit in mice with VS dopamine ablations was reflected in a failure to reach the vicinity of the water spout on individual trials. Control mice reached the water spout on most trials. Ablation mice, on the other hand, were significantly more likely to abort their travel into the arena and return to the shelter while still far from the water spout (Figure 1E-G). Importantly, the ablation effect on approach persistence was location-dependent (Figure 1G-H). While animals were far (distal) from the reward, ablation animals returned to the shelter without obtaining reward significantly more often than controls (Figure 1G right, H left). Yet as animals neared the water spout, the reward acquisition rates between groups became similar, even early in training (Figure 1G right, H right). The impairment in approach to distal reward in ablation mice persisted across training: while ablation mice improved reward acquisition at a similar pace to controls when near reward, they continued to lag behind control mice in their approach to distal reward (Figure 1H). Taken together, ablation of VS-projecting dopamine neurons induced a selective deficit in persistence towards a distal reward, while leaving engagement with a proximal reward intact. This suggests a specific role for VS dopamine in facilitating navigation toward distal reward.

### Mice with ablation of VS-projecting dopamine neurons show decreased suppression of foraging by threat

We next assessed how the introduction of threat into the environment affects the role of VS dopamine in foraging (Figure 2). If VS dopamine conveys mainly reward information, it is expected that mice with ablation of VS-projecting dopamine neurons would continue to show deficits in reward acquisition under a potential threat. Following an initial training period with no threat, a “monster” object was introduced to the distal end of the arena in some sessions (monster sessions) (Figure 2A). In monster sessions, monster movement was triggered when the mouse reached an invisible line in the arena. The monster moved forward and backward repeatedly, until the mouse retreated to the shelter. In trained mice, we ran alternating monster sessions and no-monster sessions; in the latter, as in training, the monster was absent. Consistent with a previous study^13^, the presence of the monster threat significantly decreased success in obtaining reward. Surprisingly, ablation mice were more successful than control mice at obtaining reward in the face of threat (Figure 2B-D). In other words, reward acquisition rates were less modulated by the presence of threat in the ablation mice compared to the control mice (Figure 2C-D).

**Figure 2.**
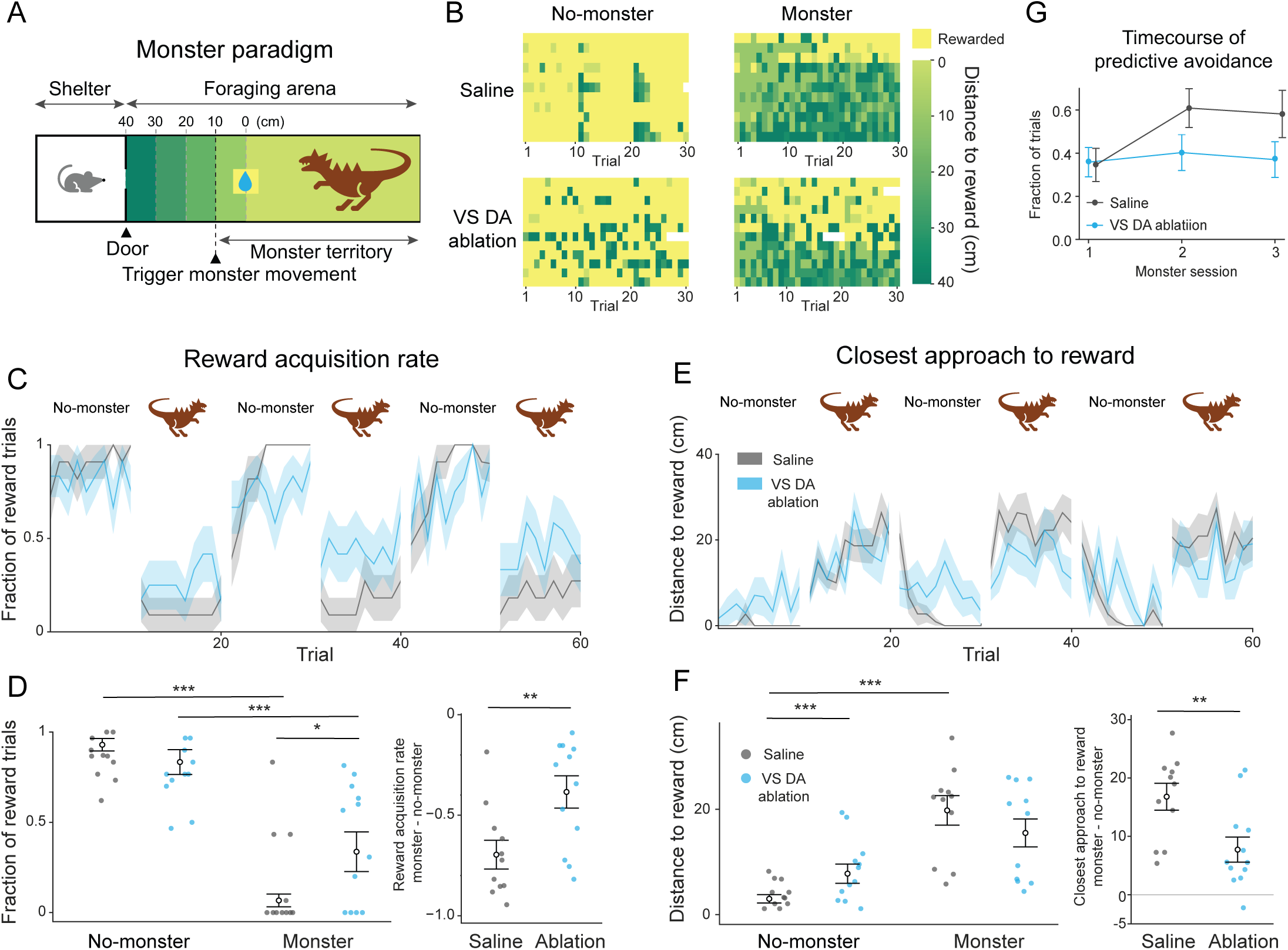
Ablation of VS-projecting dopamine neurons increases successful reward acquisition under threat. (A) Schematic of the monster task. When the mouse crossed an invisible line at 10 cm from water reward, a motorized “monster” toy was triggered to move back and forth. (B) The distance to reward at closest approach in each trial for each animal. (C) Reward acquisition rate across trials for animals with VS dopamine ablation (blue) and control (saline, gray). (D) Left, monster threat decreased reward rates in both groups. Ablation mice exhibited higher reward acquisition rate than control animals during monster sessions (Wald Z test: monster vs no-monster session, ablation, OR = 0.10, *z* = −10.8, *p* = 1.3×10⁻^26^ ; control, OR = 0.0054, *z* = −12.9, *p* = 2.2×10⁻^37^; ablation vs control, no-monster: OR = 0.37, *z* = −1.3, *p* = 0.97; monster, OR = 7.0, *z* = 2.7, *p* = 0.036). Right, threat-induced suppression of reward acquisition was attenuated by VS dopamine ablation (Welch t-test: *t*(20.9) = 2.9, *p* = 0.0086). (E) The closest approach to reward for VS dopamine ablation (blue) and control (gray) animals. (F) Threat significantly increased the distance to reward at closest approach in control animals, whereas the effects of threat decreased in ablation mice (Wald z-test: left, monster vs no- monster, control, *z* = 5.5, *p* = 2.1×10⁻^7^ ; ablation, *z* = −0.062, *p* = 1.00; ablation vs control, no- monster: *z* = 5.8, *p* = 2.3×10⁻^8^ ; monster: *z* = 0.47, *p* = 1.0; right, ablation vs control, Welch t-test: *t*(20.7) = 2.8, *p* = 0.0091). (G) Ablation mice showed significantly less development of predictive avoidance across monster sessions compared to control mice (*t*(44) = 2.7; *p* = 0.0090). Line plots, mean ± SEM. Error bars, mean ± SEM. Dots, individual animals. n = 23 animals (11 ablation, 12 control). * *p* < 0.05, ** *p* < 0.01, *** *p* < 0.001.

To further elucidate the effect of threat on foraging behavior, we next examined how far animals ventured into the arena in each trial. Consistent with effects in training, during no-monster sessions, VS dopamine ablation mice traveled significantly less distance towards the reward before returning to shelter, compared to control mice (Figure 2B, E-F). However, in the presence of threat, ablation mice traveled slightly closer to reward compared to their control counterparts. To understand this change, we investigated the impact of threat on distance traveled towards reward in each group. In monster sessions, control mice significantly decreased their distance traveled towards reward compared to no-monster sessions, staying far from the monster threat. In contrast, this approach behavior was significantly less affected by threat in ablation mice (Figure 2E-F). Indeed, while control mice, after experiencing the moving monster, began to avoid the monster early in the trial, returning to the shelter even before triggering the monster charge (“predictive avoidance”^13^), ablation mice showed significantly less development of this early avoidance of distal threat across sessions (Figure 2G). Overall, ablation mice showed reduced adjustment of foraging behavior according to threat context.

### Threat responses in TS but not VS dopamine predict avoidance behavior

The observed behavioral deficits in ablation mice suggest that VS dopamine is important for both sustaining reward-motivated behavior and appropriately suppressing such behavior to facilitate threat avoidance. Previous work has linked dopaminergic modulation in another striatal region, TS, to the appropriate suppression of reward-seeking behavior under threat^9,13^. Given our finding that VS dopamine, like TS dopamine, is important for avoidance behavior, we next compared their activity patterns.

To achieve this comparison, we simultaneously recorded the activity of dopamine axons in VS and TS using a Ca^2+^ indicator (GCaMP6f) in triple transgenic mice (DAT-Cre/Ai148/TdT) (Figure 3). We first investigated responses in each region to key events in the monster task (Figure 3D; Figure S1). Consistent with prior findings of specialized processing of reward and threat in VS and TS dopamine, respectively, monster charge evoked a significant response in TS dopamine axons but not in VS, while delivery of the water reward evoked a stronger response in VS dopamine axons than in TS.

**Figure 3.**
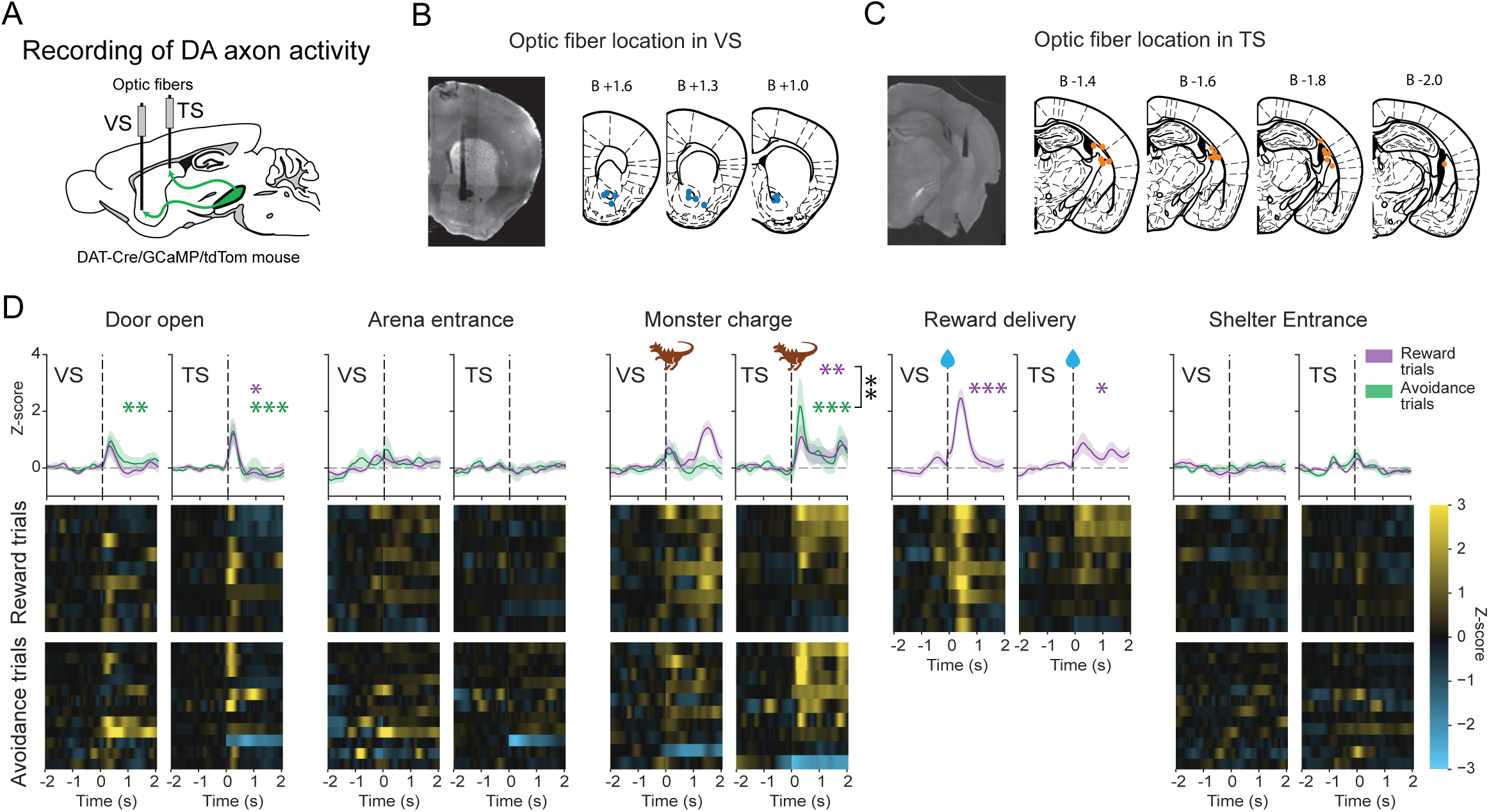
Simultaneous recordings from dopamine axons in VS and TS during foraging under threat. (A) Schematic of Ca^2+^ indicator recording from dopamine axons in VS and TS. (B, C) A representative coronal section showing an optic fiber track and TH expression (white) (left) and summary of fiber tip location across mice in VS (B) and TS (C). (D) Event-aligned signals (Z-scored) for reward acquisition trials (purple, “reward”) and avoidance trials (green, “avoidance”) in dopamine axons in VS and TS at door opening (pairwise t-test: TS, reward, *t*(192) = 2.9, *p* = 0.038; avoidance, *t*(158) = 4.0, *p* = 7.8 × 10^-4^; reward vs avoidance, *t*(248) = - 0.55, *p* = 0.58; VS, reward, *t*(235) = 2.2, *p* = 0.25; avoidance, *t*(211) = 3.3, *p* = 0.0085; reward vs avoidance, *t*(248) = −0.65, *p* = 0.52), arena entrance (one-sample t-test: TS, reward, *t*(192) = - 0.28, *p* = 1.00; avoidance, *t*(158) = −0.36, *p* = 1.00; pairwise t-test: reward vs avoidance, *t*(248) = 0.032, *p* = 0.97; one-sample t-test: VS, reward, *t*(235) = 0.94, *p* = 1.00; avoidance, *t*(211) = 1.5, *p* = 1.00; pairwise t-test: reward vs avoidance, *t*(248) = −0.37, *p* = 0.71), monster charge (one- sample t-test: TS, reward, *t*(192) = 3.3, *p* = 0.0091; avoidance, *t*(195) = 7.5, *p* = 1.9 × 10^-11^; pairwise t-test: reward vs avoidance, *t*(253) = −3.2, *p* = 0.0015; one-sample t-test: VS, reward, *t*(235) = 1.7, *p* = 0.76; avoidance, *t*(239) = 1.4, *p* = 1.00; pairwise t-test: reward vs avoidance, *t*(252) = 0.21, *p* = 0.83; TS vs VS, reward, *t*(195) = 0.65, *p* = 1.00; avoidance, *t*(199) = 3.40, *p* = 0.0040), reward delivery (one-sample t-test: TS, *t*(192) = 2.87, *p* = 0.041; VS, *t*(235) = 6.47, *p* = 5.0 × 10^-9^; pairwise t-test: TS vs VS, *t*(195) = −3.40, *p* = 7.6 × 10^-4^), shelter entrance (one-sample t-test: TS, reward, *t*(192) = 0.66, *p* = 1.00; avoidance, *t*(158) = 1.29, *p* = 1.00; pairwise t-test; reward vs avoidance, *t*(248) = −0.40, *p* = 0.69; one-sample t-test: VS, reward, *t*(235) = −0.46, *p* = 1.00; avoidance, *t*(211) = 0.20, *p* = 1.00; pairwise t-test: reward vs avoidance, *t*(248) = −0.53, *p* = 0.60). Mean ± SEM across animals. n = 12 animals. * *p* < 0.05, ** *p* < 0.01, *** *p* < 0.001.

Furthermore, the TS dopamine response to monster charge onset, but not the VS dopamine response, was predictive of animals’ subsequent decisions. Consistent with a previous study^13^, dopamine axon activity in TS at monster charge was higher on trials in which animals fled from the monster, failing to obtain reward, and lower on trials in which animals successfully acquired reward despite the monster threat (Figure 3D). The response to monster charge in VS dopamine axons, by contrast, showed no significant modulation by behavioral decision, suggesting that TS and VS dopamine are differentially associated with behavioral threat response. Critically, the response to monster charge in VS dopamine axons was not significantly negative, suggesting that this potential threat is encoded differently by VS dopamine than unpredicted physically aversive stimuli such as electric shock or air puff, which inhibit VS dopamine axons^9,24^.

Taken together, although ablation experiments showed important roles for both VS and TS dopamine in threat avoidance, event-locked responses to the potential threat differ notably between VS and TS. Monster movement evokes activation of dopamine axons in TS, and the magnitude of this response is modulated by trial outcome (either ultimate avoidance, or reward acquisition in spite of threat). VS dopamine, on the other hand, showed minimal event-locked response to monster movement, and no modulation by subsequent trial outcome. Importantly, the threat response of VS dopamine axons is not similar nor opposite to that of TS dopamine, revealing differential encoding rules in these dopamine neuron subpopulations that we could not detect in previous studies using physically aversive stimuli^8,9^.

### Reward-oriented kinematics explain VS and TS dopamine beyond generic movement encoding

Our event-locked analyses indicate distinct response patterns of VS and TS dopamine during foraging under threat. Importantly, dopamine activity in both VS and TS showed numerous transient and sustained modulations throughout the trial falling outside event-locked windows (Figure S2). This is not surprising as the behavioral repertoire is inherently broader and more complex than can be summarized by a few discrete timestamps^26^. Indeed, when we visualized dopamine activity across arena space, we observed spatially distributed modulations, with single trajectories showing strong peaks of dopamine responses at varied locations, not clearly localized to appreciable landmarks (Figure 4A-B).

**Figure 4.**
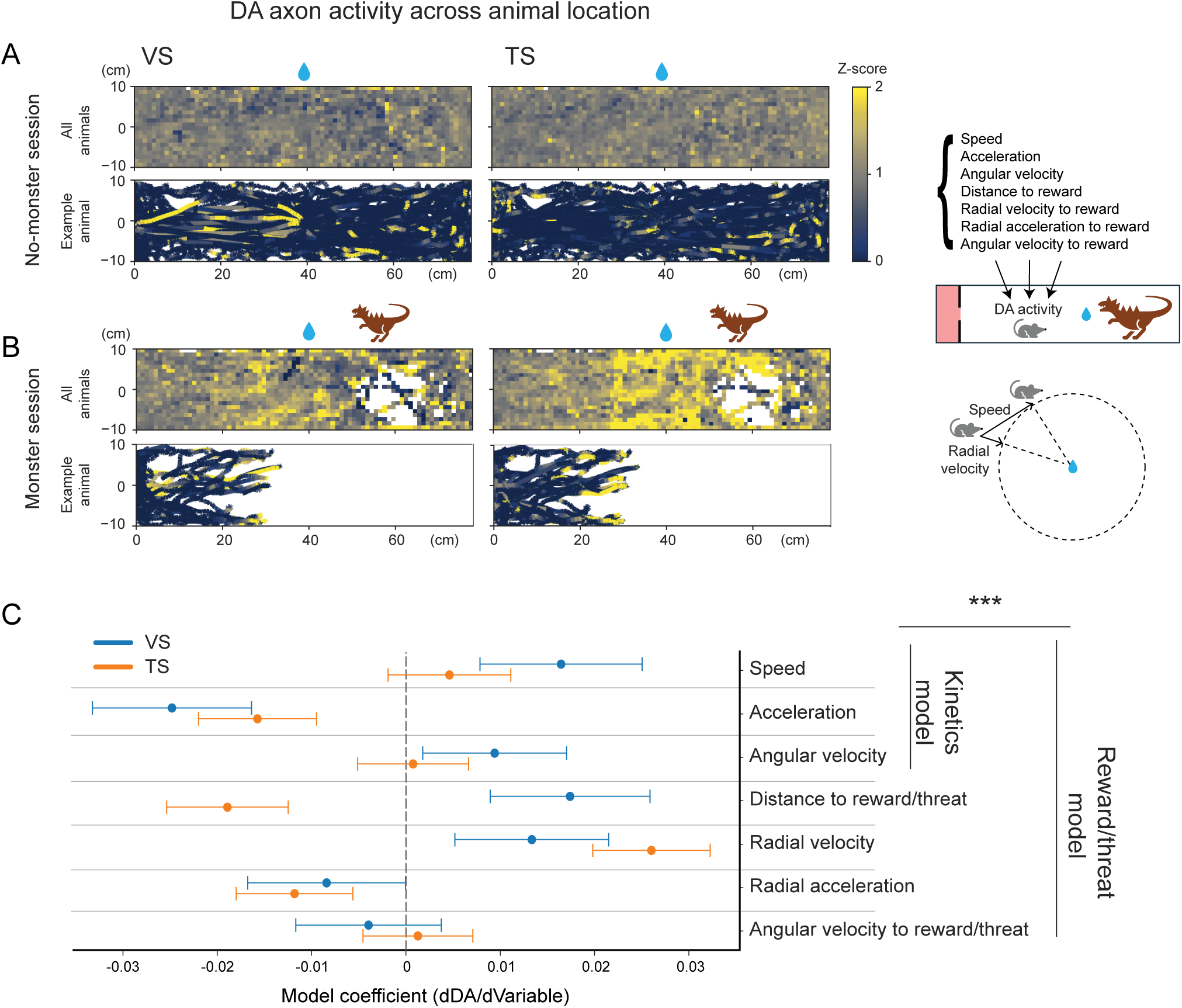
Modulation of VS and TS dopamine axon signals by kinematics relative to reward and threat. (A-B) Spatial maps showing position-binned average Ca^2+^ signals in VS (upper left) and TS (upper right) dopamine axons during the no-monster (A) and monster (B) session across all animals and trajectories (VS, lower left; TS, lower right) in an example animal. (C) Mixed-effects model of dopamine axon activity in VS (blue) and TS (orange) was constructed with animal’s movement kinematic and reward/threat related variables. Including reward/threat related variables improved explained variance for the model relative to the kinematics-only model (likelihood-ratio test: *χ²*(8) = 160.9, *p* = 2.2 × 10^-9^; AIC: 141857 vs 142002; BIC: 142031 vs 142103). The reward/threat model explained a substantial fraction of variance in dopamine axon activity (Conditional R^2^: 0.33; Marginal R^2^: 0.32). Fixed-effect estimates (changes in Z-scored DA signals/Z-scored variables ± 95% confidence intervals) for reward/threat model are shown. n = 12 animals.

Previous studies reported modulation of dopamine activity by kinematic factors such as animal’s movement speed^27–30^, acceleration^31^, and angular movement contralateral to dopamine axon location^32,33^. On the other hand, multiple studies found that dopamine dynamics not tightly locked to observable events may also encode moment-to-moment reward-related information^34–38^. We first tested whether dopamine axon activity could be accounted for by kinematic measures alone compared to reward/threat stimuli. Specifically, we fit a baseline kinematics model using core movement predictors, and then fit an extended model that added reward/threat-directed kinematic regressors (distance, radial velocity, acceleration and angular velocity toward water spout location, which is proximal to the monster threat in monster sessions) (Figure 4C). To address potential confounding by phasic reward transients in both models, reward-evoked responses were regressed out first (see Methods, Figure S3). We found that the extended model provided a better fit to dopamine activity in both VS and TS than the kinematics-only model, indicating that kinematic components relative to reward/threat explain dopamine activity beyond generic movement variables alone. Among the predictors in the extended model, radial velocity was positively correlated with dopamine axon activity both in TS and VS, whereas distance to reward/threat was positively correlated with VS dopamine and negatively correlated with TS dopamine. Importantly, radial velocity and position were not tightly correlated in our naturalistic setting as is usually the case in reward approach in a more constrained arena^34,37^, allowing us to dissociate contributions of each variable. We focused subsequent analyses on these two variables to clarify how stimulus-directed movement and spatial context shape dopamine dynamics.

### VS dopamine activity is flexibly modulated by approach or retreat kinematics under reward-threat conflict

We first examined how dopamine activity scaled with the first model-identified predictor, radial velocity toward reward/threat (Figure 5, S4). This is a signed kinematic measure defined along the shelter to water spout axis (see Figure 4), with positive values indicating movement toward the reward/threat, and negative values indicating movement toward the shelter. Previous studies found striatal dopamine signals to exhibit sustained ramping dynamics as animals navigate toward reward, mirroring progressive changes in expected value^27^. Subsequent work demonstrated that this activity covaries with ongoing visual scene movement speed, reflecting a temporal-difference (TD) reward prediction error (RPE) as animals virtually move along an expected value trajectory^34^. To investigate whether VS and TS dopamine similarly reflect evaluation of an expected value (or threat) trajectory, we isolated epochs of approach to the reward/threat (see Methods), which were dominated by positive radial velocities, and retreat epochs, which were dominated by negative radial velocities (Figure 5A). We asked how radial velocity and animal location related to dopamine activity during approach and retreat epochs in monster sessions. To examine the tripartite relationship between radial velocity toward reward/threat, animal location in the arena, and dopamine axon activity, we plotted individual animal trajectories across the arena during periods of high dopamine axon activity, and visualized radial velocity throughout the trajectory (Figure 5A). Interestingly, compared to TS dopamine (Figure 5A bottom), trajectories characterized by high VS dopamine activity corresponded to high velocities and were relatively localized along the center of the track, suggesting straight runs towards reward and away from the monster (Figure 5A middle). Consistently, we noticed that VS dopamine activity tended to be high when the radial velocity is high or peaked in an example trajectory (Figure 5B top).

**Figure 5.**
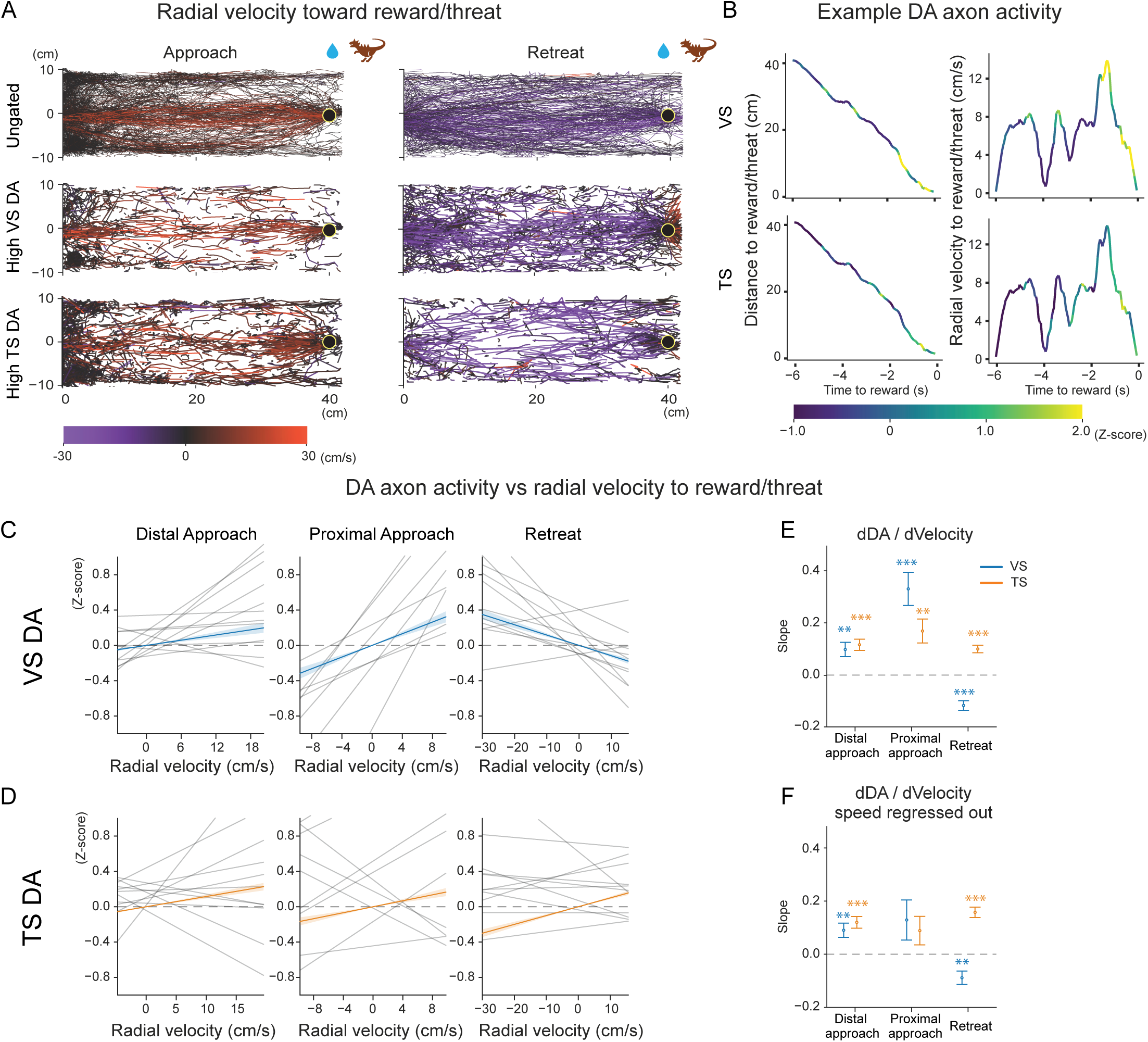
**VS dopamine axon activity is flexibly modulated by animal’s relative movement to reward and threat in a behavioral context-dependent manner**. (A) Top, example animal trajectories during approach (left) and retreat (right) epochs, color-coded by radial velocity relative to reward/threat position (positive values indicate movement toward reward/threat; negative values indicate movement toward the shelter). Middle, trajectories from time points with high VS dopamine axon activity (Z > 1) during approach and retreat. Bottom, trajectories from time points with high TS dopamine axon activity (Z > 1) during approach and retreat. (B) Example of how VS (top) and TS (bottom) axonal activity varies with distance (left) and radial velocity (right) to reward/threat in an exemplar mouse in a single trial. (C, D) Correlation between animal radial velocity toward reward/threat and dopamine axon activity in VS (C) and TS (D). Thin black lines, per-animal regression slopes. Colored lines (blue, VS; orange, TS), the population-level mixed-effects regression slope ± SEM. (E) Estimated slopes for VS (blue) and TS (orange) dopamine axon activity vs radial velocity (dDA/ dVelocity) across behavioral epochs (one sample Wald z-test: VS: distal approach, *z* = 3.560, *p* = 0.0037; proximal approach, *z* = 5.191, *p* = 2.1 × 10^-6^; retreat, *z* = −6.279, *p* = 3.4 × 10^-9^; TS: distal approach, *z* = 5.421, *p* = 5.9 × 10^-7^; proximal approach, *z* = 3.676, *p* = 0.0024; retreat, *z* = 6.941, *p* = 3.9 × 10^-11^). (F) Estimated slopes of dopamine axon activity vs radial velocity (dDA/dVelocity), with speed regressed out, across behavioral epochs (one sample Wald z-test: VS: distal approach, *z* = 3.3, *p* = 0.0079; proximal approach, *z* = 1.7, *p* = 0.88; retreat, *z* = −3.5, *p* = 0.0046; TS: distal approach, *z* = 5.7, *p* = 7.9 × 10^-8^; proximal approach, *z* = 1.649, *p* = 0.99; retreat, *z* = 8.047, *p* = 8.5 × 10^-15^).Open circles indicate the mean; error bars indicate ± SEM. n = 12 animals. * p < 0.05, *** p < 0.0001.

To confirm this impression of distinct relationships between radial velocity and VS vs TS dopamine, we used mixed-effects modeling to quantify the linear correlations between dopamine activity and radial velocity toward reward/threat. Consistent with our observations of raw data (Figure 5A-B), we found a marked dissociation between VS and TS dopamine. In VS, dopamine axon activity was strongly correlated with radial velocity during both approach and retreat behavior. During approach, activity in VS increased with increasing radial velocity toward reward/threat, and this positive slope was significant both when animals were distal and proximal to reward/threat (Figure 5C, E). After retreat onset, the sign of the relationship reversed, such that dopamine axon activity in VS was higher during faster retreat to the shelter, resulting in a significantly negative slope relating VS dopamine to radial velocity (Figure 5C, E). Importantly, this approach vs retreat-dependent sign reversal in the relationship between VS dopamine and radial velocity was specific to sessions with the monster threat present. In the no-monster sessions, dopamine axon activity in VS remained positively correlated with velocity toward the reward during approach but did not exhibit a negative correlation during retreat (Figure S4C, E-F). Thus, the flip in the relationship between VS dopamine axon activity and the animal’s radial velocity toward reward is not simply due to the animal’s movement direction, but rather elicited by the presence of threat. Consistent with this, repeating the radial-velocity analyses after regressing out unsigned speed recapitulated the original tendencies (Figure 5F), further supporting the conclusion that these radial velocity-linked dopamine effects are not reducible to locomotor speed alone. These results suggest that VS dopamine signals the global motivational landscape, influenced by both environmental (presence or absence of threat) and behavioral (approach or retreat) context. When animals approach reward, the motivational gradient tracked by VS dopamine is directed towards reward, with or without the monster threat present. When animals decide to retreat under threat – a change in the behavioral context – the gradient flips away from the reward and threat location, such that faster retreats toward the shelter are accompanied by elevated dopamine. However, when animals decide to retreat under no threat, there is no obvious gradient developed (Figure S4), so the switch depends on both environmental (threat) and behavioral (retreat) context.

In contrast to VS dopamine, which showed context-dependent correlations with radial velocity, TS dopamine exhibited a consistently positive relationship with radial velocity during both approach and retreat in monster sessions (Figure 5D-F), implying a consistent threat gradient towards the threat location.

Together, these results suggest that VS dopamine mirrors the animal’s velocity towards a location with higher value, with the value landscape set by the current environmental and behavioral context. Overall, VS dopamine, but not TS dopamine, shows a context-dependent relationship with an animal’s movement, correlating with velocity towards an animal’s current goal.

### VS dopamine as a behavioral context-dependent TD RPE under reward-threat conflict

The dynamic, context-dependent relationship between VS dopamine and radial velocity suggests that VS-projecting dopamine neurons are more activated when animals move toward a location with higher value relative to the animal’s current location in space (i.e. toward water reward during approach, and away from a threat during retreat). These modulation patterns fit precisely with the idea of VS dopamine encoding a TD-RPE of value over space (Figure 6A). In TS dopamine, on the other hand, a positive correlation with velocity toward the reward/threat location was seen, which could reflect these dopamine neurons computing an analogous TD prediction error of threat over space. However, a positive correlation was also observed in no- monster sessions, so this case is more complex (Figure S4). To assess whether dopamine axon activity in VS and TS reflects a TD-RPE of spatial value (or threat), we integrated dopamine signals across time (see Methods, Figure 6B), then visualized the accumulated signals across space during approach vs retreat in our two environmental contexts (with or without monster threat) (Figure 6C-F, S5). We reasoned that if dopamine axon activity reflects a TD-RPE, a derivative-like transformation of value, its temporal integral should approximate the underlying value map in each context.

**Figure 6.**
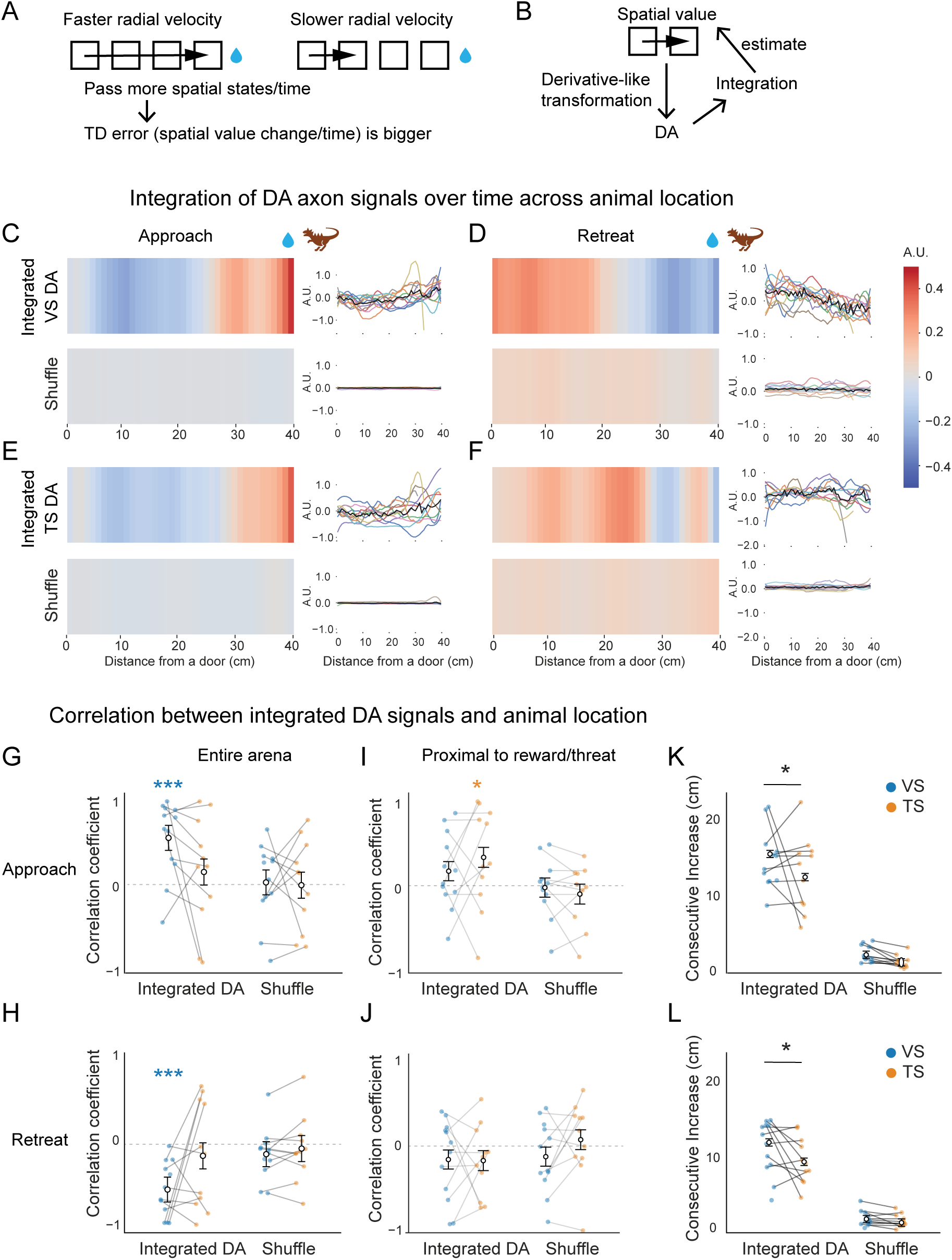
Integrated VS dopamine axon signals form spatial gradients toward reward/threat, that reverses upon retreat. (A) Relationship between radial velocity to reward and TD RPE. (B) If dopamine signals approximate TD RPE of spatial value, integration of dopamine signals may estimate spatial value. (C-F) Integrated dopamine axon signals in VS (C, D) and TS (E, F) plotted as a function of the position in the arena during approach (C, E) and retreat (D, F). Heatmaps (left) show the mean position-binned integrated signals for the observed data (top) and a within-trial circularly permuted control (bottom, “shuffle”). Line plots (right) show signal traces in each animal across the arena (colored lines) with the group mean overlaid (black). (G-H) Monotonicity of the gradient during approach (G) and during retreat (H) was tested with Spearman correlation. Only the mean Spearman correlation of the raw VS signals, but not TS signals, differed significantly from zero (pairwise t-test: approach, VS, raw, *t*(168)=3.7, *p*=0.0010; shuffle, *t*(168)=0.16, *p*=1.00; TS, raw, *t*(168)=0.96, *p*=1.00; shuffle *t*(168)=−0.051, *p*=1.00; retreat, VS, raw, *t*(168)=−3.6, *p*=0.0015; shuffle *t*(168)=−0.793, *p*=1.00; TS, raw, *t*(168)=−0.88, *p*=1.00; shuffle, t(168)=−0.32, *p*=1.00). (I-J) The same analysis as G-H but using the data proximal to reward/threat was assessed. Only the mean Spearman correlation for the TS signals differed significantly from zero during approach (pairwise t-test: approach, TS original: *t*(162) = 2.8, *p* = 0.018; VS original: *t*(160) = 1.5, *p* = 0.50; TS shuffle: t(162) = −1.5, *p* = 0.47; VS shuffle: *t*(160) = −0.47, *p* = 1.00; retreat, TS original: *t*(162) = −1.4, *p* = 0.54; VS original: *t*(160) = −1.4, *p* = 0.60; TS shuffle: *t*(162) = 0.22, *p* = 1.00; VS shuffle: *t*(160) = −1.0, *p* = 1.00). (K-L) Consecutively increasing spatial bins during approach (K) and during retreat (L) are significantly longer in VS than in TS (Wald z-test: approach, VS vs TS, raw data, *z*=−5.9, *p* = 2.3 × 10^-8^; shuffle control, *z*=−1.9, *p*=0.33; raw vs shuffle, VS, *z*=26.0, *p* = 1.2 × 10^-148^; TS, *z*=21.5, *p* = 8.7 × 10^-102^; retreat, VS vs TS, raw, *z*=−4.8, *p* = 8.8 × 10^-6^; shuffle, *z*=−0.89, *p*=1.00; raw vs shuffle, VS, *z*=19.3, *p* = 2.3 × 10^-82^; TS, *z*=14.9, *p* = 6.3 × 10^-50^). Error bars, mean ± SEM. Dots, individual animals. n = 12 animals. * *p* < 0.05, *** *p* < 0.001.

We observed distinct patterns in the integrated signals during approach vs retreat epochs, and between integrated VS vs TS dopamine axon signals. In VS, the integrated signals increased progressively as animals advanced toward reward/threat during approach (Figure 6C), consistent with steadily increasing value. Importantly, this spatial pattern was not present in shuffle controls (see Methods), arguing against a trivial explanation based on arena occupancy or analysis artifacts (Figure 6C). Strikingly, the integrated signals showed a corresponding ramp in the opposite direction during retreat, rising as animals moved back toward the shelter (Figure 6D).

This bidirectional ramping is consistent with VS dopamine flexibly evaluating the particular value landscape relevant to the animal’s current goal, with value increasing towards reward during approach, and away from threat or towards safety during retreat. Interestingly, we did not observe clear gradients in no-monster sessions (Figure S5A-D), suggesting that value maps were more complicated or animal movement was not as acutely goal-directed in no-monster sessions. To assess how consistently the integrated VS dopamine signals were monotonically modulated by an animal’s location in the arena, we examined correlation coefficients between the signals and distance from reward/threat. We found consistent monotonicity in VS during both approach (Figure 6G) and retreat (Figure 6H), supporting the idea that VS dopamine reflects a behavioral context-dependent TD-RPE of spatial value across the foraging arena.

In contrast to the continuous ramps towards a goal observed in integrated VS dopamine signals, integrated TS dopamine signals did not show clear monotonic changes across the arena (Figure 6E-H). However, despite this lack of overall monotonicity of integrated signals (Figure 6G-H), the average temporal integration in TS dopamine signals did show a ramp-like pattern during approach, especially near the threat location (Figure 6E). One possibility is that the ramp in TS dopamine activity during approach is evoked by direct sensory experience, present only when the monster is visible. Indeed, the ramps calculated from TS dopamine show a sharp upturn at 30cm (Figure 6E), the location at which the monster threat is activated to begin charging, and the sensory stimulus is thus magnified. Accordingly, when the animal was proximate to the charging monster, TS dopamine developed significant monotonicity during approach (Figure 6I), despite not showing monotonicity across the full arena (Figure 6G). Furthermore, the distance over which the integrated signal increased was significantly shorter in TS than in VS (Figure 6K, L), suggesting that TS dopamine preferentially evaluates local information proximal to threat, whereas VS dopamine evaluates properties of the environment more globally.

### TS dopamine activity is shaped by threat proximity and direction

Recalling that distance to reward/threat emerged as another key predictor of dopamine activity in our regression model (Figure 4), we next considered the relationship between dopamine and position relative to reward/threat (Figure 7, S6, S7). One possible explanation for the observed modulation of dopamine activity in TS by distance to reward/threat, independent of the current goal, is that activity of TS dopamine neurons is reflective of immediate sensory stimuli, rather than underlying structure of the environment. In other words, TS dopamine may be modulated by egocentric sensory experience relative to the animal, rather than by the allocentric space through which the animal moves.

**Figure 7.**
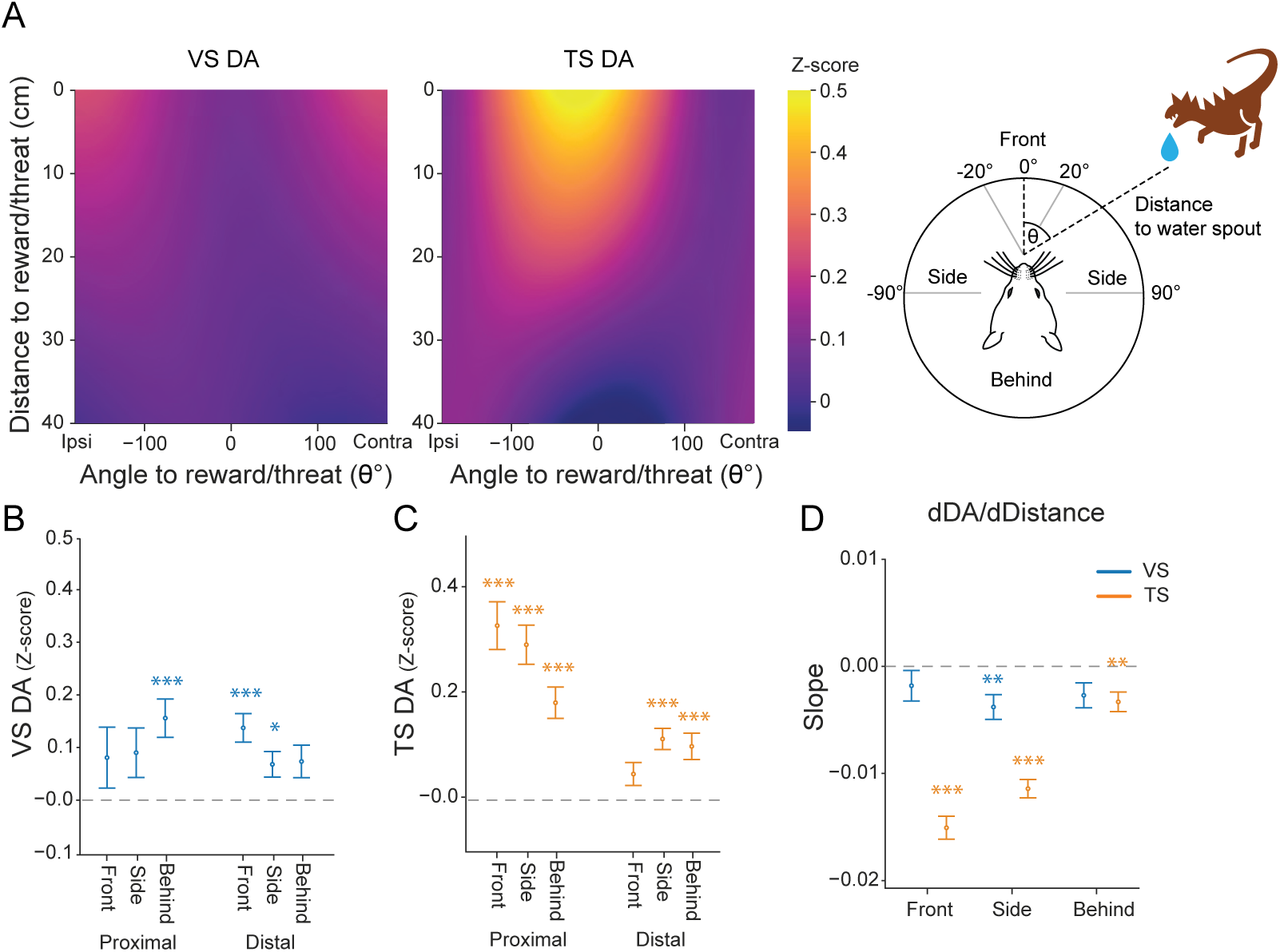
**TS dopamine axon activity is modulated by threat proximity and direction. (**A) Two-dimensional maps of z-scored dopamine axon activity as a function of head angle relative to and distance to the reward/threat for VS (Left; pairwise t-test ipsi vs contra: *t*(2036) = 1.02, *p*=0.31) and TS (Middle; pairwise t-test ipsi vs contra: *t*(536) = −0.50, *p*=0.62). Color indicates model-estimated mean z-scored fluorescence across samples within each angle-distance bin. Positive and negative angles indicate the side contralateral and ipsilateral to the recording side, respectively. (B-C) Mean dopamine axon activity in VS (B) and TS (C) as a function of proximity to the reward/threat (proximal, distal) and orientation relative to the reward/threat (front, side, behind) (one sample Wald z-test: VS, proximal, front, *z* = 1.4, *p* = 0.98; side, *z* = 1.9, *p* = 0.33; behind, *z* = 4.3, *p* = 1.2 × 10^-4^; distal, front, *z* = 5.1, *p* = 2.4 × 10^-4^; side, *z* = 2.8, *p* = 0.031; behind, *z* = 2.4, p = 0.11; TS, proximal, front, *z* = 7.3, *p* = 1.5 × 10^-12^; side, *z* = 7.9, *p* = 1.0 × 10^-14^; behind, *z* = 6.2, *p* = 3.0 × 10^-9^; distal, front, *z* = 2.3, *p* = 0.13; side, *z* = 5.8, *p* = 2.1 × 10^-8^; behind, *z* = 4.1, *p* = 2.7 × 10^-4^). (D) Slope of dopamine axon activity with respect to distance to reward/threat (dZ/dDistance) within each orientation category (front, side, behind) for VS (blue) and TS (orange). Negative slopes indicate higher dopamine at positions closer to reward/threat. TS showed markedly steeper negative distance slopes than VS across orientation categories, particularly when animals were directly facing reward/threat (Wald z-test: VS, front, *z* = −1.3, *p* = 1.0; side, *z* = −3.3, *p* = 0.0064; behind, *z* = −2.3, *p* = 0.12; TS, front, *z* = −14.1, *p* = 2.8 × 10^-44^; side, *z* = −13.2, *p* = 5.4 × 10^-39^; behind, *z* = −3.6, *p* = 0.0018; VS-TS, front, *z* = −7.5, *p* = 8.3 × 10^-14^; side, *z* = −5.3, *p* = 1.3 × 10^-7^; behind, *z* = −0.41, *p* = 0.68). Error bars, mean ± SEM. n = 12 animals. * *p* < 0.05, ** *p* < 0.01, *** *p* < 0.001.

To test this idea, we investigated whether dopamine axon activity in TS depends on the animal’s proximity to reward/threat and head orientation relative to these stimuli (Figure 7). We plotted TS and VS dopamine activity as a function of distance to and angle to reward/threat, which revealed strong modulation by animal position relative to reward/threat in TS dopamine, and comparatively minimal modulation in VS dopamine (Figure 7A, S7E). The relationship between TS dopamine activity and distance to reward/threat was notably modulated by head orientation relative to these stimuli; TS dopamine activity was highest when animals were close to and facing reward/threat, but the modulation by distance was substantially attenuated when the reward/threat were behind the animal. To quantify the impact of orientation and distance to the reward/threat on TS dopamine activity, we categorized each animal’s orientation into three sectors: front (within the animal’s binocular zone, ±20°^39,40^), side (±20 to ±90°), and behind (≥ ±90°) (Figure 7A). We then used mixed-effects models to estimate mean dopamine activity within each orientation-distance bin (Figure 7B-C). This analysis revealed pronounced spatial tuning in TS dopamine: TS dopamine axon responses were highest when animals were proximal to and facing towards the reward/threat, whereas responses were reduced when the reward/threat was behind the animal, or at far distances (Figure 7A, C). By contrast, VS dopamine showed smaller effects of distance or angle (Figure 7B). Modeling dopamine as a continuous function of distance within each orientation category showed that VS exhibited little dependence on distance, whereas TS displayed substantially steeper, orientation-dependent distance slopes (Figure 7D), indicating stronger proximity coding when the reward/threat lies in front of the animal and weaker proximity dependence when it is to the side or behind. When our mixed-effects model was fit in no-monster sessions, in contrast, TS dopamine showed largely attenuated modulation by distance and orientation to the same location, compared to monster sessions (Figure S6). Thus the observed modulation appears to be mainly due to the presence of the monster threat. Notably, we did not observe lateralization in TS dopamine activity, which responded similarly to stimuli in contra- vs ipsi-lateral locations (Figure 7A, S6A).

Taken together, these results reveal TS dopamine to be especially sensitive to local sensory stimuli, tightly encoding an animal’s spatial relationship to the monster threat. The TS dopamine response to monster charge was associated with whether animals ultimately persisted in approach or retreated (Figure 3), which suggests this local sensory signal may reflect a threat estimate to inform subsequent behavior. In contrast, VS dopamine showed comparatively rudimentary sensory encoding, with activity instead consistent with a TD-RPE that flexibly supports the behavior (approach, retreat) most advantageous within the current global context.

## Discussion

In this study we utilized a naturalistic foraging paradigm to identify collaborative roles of VS and TS dopamine to threat avoidance. Ablation of VS-projecting dopamine neurons reduced persistence to pursue distal reward in the absence of threat, yet also blunted adjustment of foraging behavior upon introduction of threat. Recordings demonstrated that VS dopamine tracked an animal’s movement relative to reward/threat in a behavioral context-dependent manner: while VS dopamine activity was correlated with radial velocity toward reward during reward approach behavior, it became correlated with radial velocity away from threat once an animal retreated toward shelter. While it is challenging to estimate value functions in a naturalistic environment, modulation of VS dopamine activity by radial velocity (the derivative of distance from reward) matches the derivative-like transformation in TD RPE, which is calculated as a discrepancy between reward prediction (value) at present and value at a previous moment ^41^. In contrast, we found TS dopamine activity to be correlated with the animal’s proximity and orientation to a threat, indicating a tight relationship with direct sensory experience. Together, these findings suggest a division of labor in which VS dopamine preferentially supports context-dependent navigation towards valued locations across space, while TS dopamine facilitates detection of and immediate retreat from threat ^13^, complementary roles which together enable effective avoidance of threat (Figure 8).

**Figure 8.**
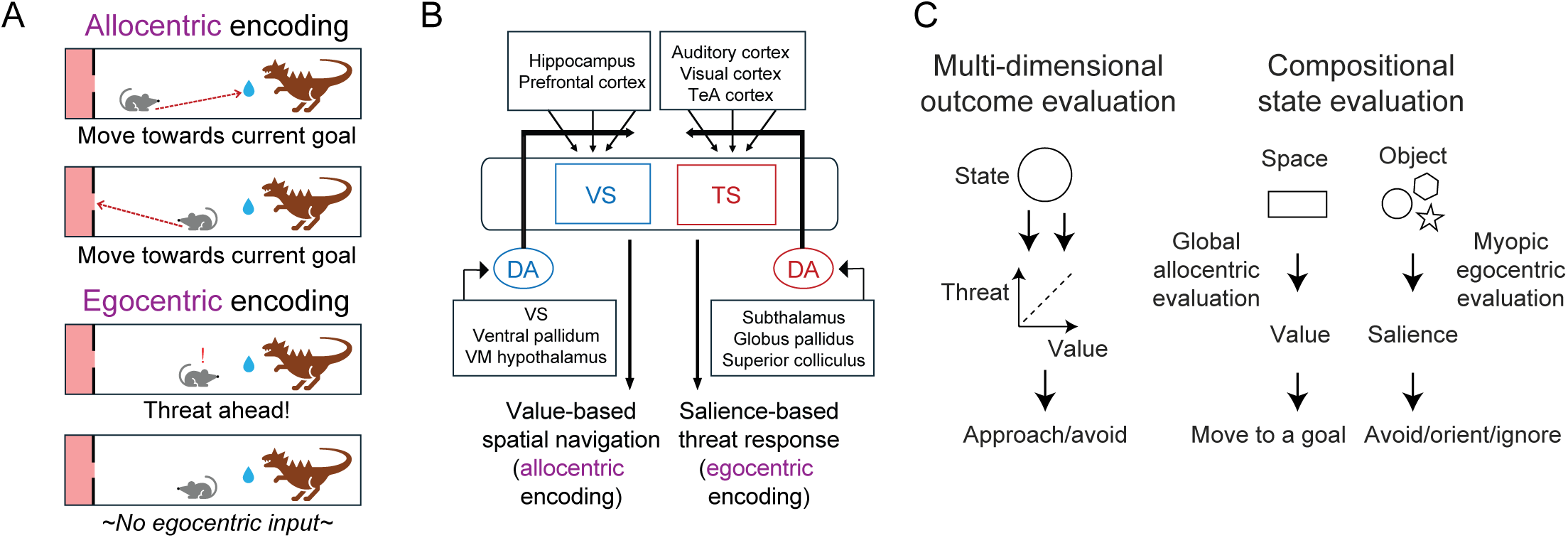
Proposed specialization of VS and TS dopamine under reward-threat conflict. (A) Schematic indicating the distinction between allocentric and egocentric encoding in the monster task, potentially supported by VS and TS. (B) Proposed model for an anatomy-imposed specialization of VS and TS for allocentric and egocentric encoding, respectively. (C) Left, previous models for dopamine diversity in TS and other striatal regions focused on outcome-specific evaluation such as flexible vs stable value, or reward vs motivational salience, threat, stimulus salience, sensory stimulus or action. Right, we propose that VS and TS dopamine evaluates a global and local state, allowing synergistic decision-making.

### VS dopamine signals a TD error of spatial value to navigate the reward-threat conflict

The VS dopamine pattern we observe may be interpreted through a TD framework, in which dopamine signals reflect a prediction error tracking moment-to-moment changes in expected future return. A previous study with head-fixed animals found that dopamine activity in VS was best fit with a first-order derivative of spatial value, consistent with TD RPE^34^. The current study extends this idea to a naturalistic foraging setting, finding that TD RPE coding in VS dopamine persists in our mixed valence paradigm.

Our proposal that VS dopamine facilitates threat avoidance by reflecting a value gradient towards the shelter builds on prior observations that dopamine neurons can signal “safety” or successful avoidance of aversive stimuli^19,22,42^. Our study reveals that dopamine activity in VS during threat avoidance can be explained as a simple TD RPE computation: during avoidance, VS dopamine neurons reflect the gradient of value away from threat, just as they reflect the gradient of value towards reward^43^.

A critical advance in our paradigm, compared to previous studies of the role of VS dopamine in reward approach and threat avoidance, is that animals in the monster paradigm can freely move forward towards reward or retreat towards the shelter at will, rather than these epochs being experimenter-prescribed. In such a naturalistic situation, we found that the TD RPE signaled by VS dopamine seems to flexibly reflect varying value landscapes determined by environmental and behavioral context. Specifically, when an animal faces both reward and threat at the same time, the activity of VS-projecting dopamine neurons was correlated with radial velocity towards a reward during approach, and with radial velocity away from a threat during retreat, apart from animal’s locomotion speed. This finding suggests that VS dopamine flexibly evaluates the environment, depending on an animal’s current goal. Flexible remapping of environmental value according to behavioral context may be triggered by upstream brain areas such as hippocampus and retrosplenial cortex^44,45^, with VS dopamine either helping to implement this remapping directly or simply reflecting a pre-formed map, to encourage a behavioral shift from reward-directed approach to retreat when a potential threat is detected.

Consistent with recording results, manipulation experiments indicate that animals with ablation of VS-projecting dopamine neurons were impaired in sustaining behavioral programs across both safe (no monster) and potentially dangerous (monster) sessions. They were specifically impaired in reliably traveling far enough into the arena to acquire distal reward, but not in acquiring proximal reward. They were also impaired in avoidance behavior, showing reduced suppression of foraging due to the presence of threat. These results demonstrate a role of VS dopamine in sustaining behavior towards reward and away from threat. Thus, the VS dopamine system does not appear to promote reward approach or threat avoidance specifically but rather allows animals to behave predictively by facilitating navigation towards a better place, whether that be movement towards more positive value or away from negative value.

### Collaboration of VS- and TS-projecting dopamine neurons under reward-threat conflict

While many studies have now reported diverse signals and functions of dopamine neurons, it is not well understood how diverse dopamine subpopulations work together. A large body of evidence implicates dopamine in VS as a reward-approach or invigoration system^35,46,47^, yet a parallel line of work has long identified its role in avoidance or learning from punishments^16–18,48–50^. On the other hand, recent work has discovered a specialized function for TS-projecting dopamine neurons in mediating threat avoidance^9,10,12,13,51^. This raised the question of what roles VS and TS dopamine play in mediating avoidance behavior, and whether these roles are complementary or opponent.

Our observation that VS dopamine plays a role in navigation to a distal reward/shelter, but not in sensory-driven reactions to a proximal reward/threat, is consistent with previous studies showing that dopamine in VS plays a role especially when sequential or sustained actions are needed^4,52^. Similarly, prior work revealed a role for VS dopamine in active avoidance^53–55^, but not in Pavlovian fear conditioning^53^. These studies suggest that VS dopamine may facilitate not only geometric, but also cognitive navigation. Mechanistically, the idea of VS dopamine facilitating value-guided navigation is consistent with the idea that dopamine in VS uniquely supports sequential conditioning^56^, a hallmark of TD learning, whereas dopamine in other striatal areas may not^4,54^. While multiple brain areas may redundantly signal positive or negative event values to facilitate association with a proximal sensory cue, VS dopamine may be crucial for animals to extend these associations across geometric and/or cognitive space, and to navigate these spaces in various task settings (Figure 8A).

In contrast to our observation that VS dopamine impacts the ability to predictively avoid potential threat but leaves initial reactive escape from proximal threat intact, a previous study^13^ found that TS dopamine ablation impairs both predictive avoidance of distal threat and reactive avoidance of proximal threat, dramatically decreasing reactive avoidance from the very first trial. Notably, TS dopamine activity was highly modulated by animal orientation towards and proximity to the monster threat. Therefore, we propose that TS dopamine is involved in creating threat predictions not across space, but mapped onto sensory experiences directly in view of the animal (Figure 8A).

The disparate modulation of VS and TS dopamine we observe during foraging under threat may reflect the distinct information each subpopulation receives (Figure 8B). Anatomically, VS receives dense input from prefrontal areas and the hippocampal navigation system^57^, a brain area that TS may not have direct access to^58^. In contrast, TS receives dense inputs from primary and higher sensory cortices including object areas such as temporal association cortex, and di- synaptically outputs to superior colliculus, an evolutionarily conserved region critical for orienting towards or away from salient stimuli^15,59^. Similarly, dopamine neurons projecting to VS and TS were also shown to receive distinct patterns of inputs^60^. Notably, this anatomical study showed that whereas most dopamine neurons are under control of VS with heavy monosynaptic inputs, TS-projecting dopamine neurons are uniquely segregated from such an information flow from VS. Our results may reflect VS- and TS-projecting dopamine neurons using the distinct state information they receive to compute TD errors supporting global and local evaluation, respectively. Without either system, animals are at a disadvantage: without VS dopamine, they appear to make myopic decisions, without utilizing the broader motivational landscape, while without TS dopamine they cannot properly react to and rapidly learn about a salient stimulus in view.

Our conclusion that avoidance behavior requires two separate dopamine systems, VS and TS dopamine, recalls the concept of compositional learning^61,62^. This theory posits that for navigation, spatial and cognitive maps are composed by hippocampus from multiple cortical representations (“building blocks”)^61^. This compositional representation allows greater generalization in new environments, both in supervised learning^61^ and in reinforcement learning models with just two representations, egocentric and allocentric^62^. The latter study proposes that although egocentric systems were previously thought to be either primitive pre-processing steps to feed into allocentric systems or to support specific actions such as skill and habits, egocentric and allocentric state evaluation systems work together to support navigation through complex environments. This scenario directly relates to our current findings, in which VS dopamine appears to evaluate spatial value in an allocentric manner, while TS dopamine evaluates sensory experience in an egocentric fashion. Taking our findings together, we propose that in the brain, TS functions as an egocentric sensory system which actively collaborates with a VS allocentric system to facilitate the complex, multistep behaviors needed to survive in natural environments (Figure 8). For avoidance of threat, TS may support quick, egocentric movements away from a salient stimulus (threat), while VS may support allocentric navigation in the direction of the better location in the environment, allowing for a more durable escape after the initial, TS-guided reaction. TS and VS dopamine signals, respectively, would train these egocentric and allocentric behavioral programs. Notably, both dopamine in TS^13^ and dopamine in VS (present work) contribute to predictive avoidance, suggesting collaboration of both local and global evaluation, especially when the situation is new (e.g. previously encountered monster threat is observed distally) as has been proposed^62^.

### Relation to previous models for diversity of dopamine responses to reward and punishment

One prominent hypothesis for dopamine’s varied responses to rewarding and aversive stimuli is that dopamine supports reinforcement learning by sending a bidirectional teaching signal as a “common currency” of net value. Recent findings including this study instead revealed the existence of diverse dopamine signals. Prior works have proposed that different dopamine neurons may evaluate different outcome information: reward value vs motivational salience^63^, flexible vs stable value^64^, reward vs threat or stimulus salience^8,9^, reward vs sensory stimulus^65^, and reward vs action^33^. Such specialized evaluation streams may allow action selection to draw flexibly on multi-dimensional outcome information^6,66^ (Figure 8C left). In addition to specialized outcome information, our findings suggest that each dopamine system may evaluate states from different perspectives such as space or object (Figure 8C right). This concept fits the anatomical reality of corticostriatal circuitry, in which each dopamine evaluation system and each striatal subregion have access to a limited subset of state information^58,60^. We propose that diversity in striatal dopamine arises not just in the service of utilizing vectorized outcome prediction errors^66–70^, but also to evaluate separate world models in support of compositional decision-making.

In summary, we found that dopamine in VS and TS collaboratively facilitates avoidance of a potential threat. Based on findings from this and prior studies, we propose that VS dopamine may facilitate allocentric value-based navigation across space while TS dopamine facilitates egocentric sensory-driven orientation/avoidance, with the two systems together allowing for adaptive avoidance behaviors. These differential roles potentially stem from distinct evolutionary pressures: the inevitably navigational nature of foraging vs the inevitably egocentric sensory-driven threat avoidance. These behavioral motifs may originally have been regulated in largely segregated brain areas before being combined to support complex behavior such as foraging in mammals.

## Methods

### Animals

A total of 35 female (n=18) and male (n=17) mice, 2-6 months old, were used. For fiber fluorometry, we used a triple transgenic mice (n=12, 8 females, 4 males) (DAT-Cre+/-; GCaMP6f+/-; tdTomato+/-), heterogygous for dopamine transporter (DAT)-Cre (B6.SJL-*Slc6a3^tm1.1(cre)Bkmn^*/J, The Jackson Laboratory #006660^71^), Cre-dependent GCaMP6f (B6.Cg- *Igs7^tm1^*^48^*^.1(tetO-GCaMP6f,CAG-tTA2)Hze^*/J or Ai148D, Jackson Laboratory #030328^72^, and Cre-dependent tdTomato (B6.Cg-*Gt(ROSA)26Sor^tm^*^14^*^(CAG-tdTomato)Hze^*/J, The Jackson Laboratory #007914^73^). For ablation experiments, we used C57BL/6J mice (saline: n = 11, 6 males/5 females; ablation: n = 12, 7 males/5 females; The Jackson Laboratory #000664). All animals were housed on a 12 hour dark/12 hour light cycle (dark from 9:00 to 21:00), and experiments were performed during the dark cycle. All procedures were performed in accordance with the National Institutes of Health Guide for the Care and Use of Laboratory Animals and approved by the Harvard Animal Care and Use Committee.

### Monster paradigm apparatus

The monster paradigm apparatus was constructed as described previously^13^ with slight modifications. The apparatus consisted of a long rectangular box (90 cm x 20 cm, 30cm tall) of white acrylic with a clear ceiling. It was divided into two compartments with a wall (and inset door) made of red acrylic. The smaller (12cm-long) compartment, referred to as the “shelter” was covered with a lid of the same red acrylic as the wall separating itself from the larger compartment. The larger (78cm-long) compartment, referred to as the “foraging arena” had a white floor and clear acrylic ceiling. The difference in ceiling color allowed the shelter to be darker (∼70% darker) than the arena. Across the shelter and arena, the ceiling had a narrow slit (∼5cm) in the center to allow the patch cord attached to mice in fluorometry experiments to follow the animal’s movement. The apparatus was automated to minimize experimenter involvement, controlled by Teensy 3.2 and Python software. The door was opened and closed with a servo motor. Infrared (IR) break beam sensors were installed at key points along the wall to detect animal movements: a sensor 1cm outside the door detected movements into the arena; a beam break at a sensor 30cm from the door, immediately before the water spout, triggered monster movement; and a beam 1cm inside the shelter detected mouse exit from the arena, triggering the door to close. A water spout was presented at the trial spout through a small hole in the floor at 40cm from the door, and licking was detected using a touch sensor. At the end of the trial, the spout was withdrawn back through the floor. The animal behavior was recorded using a USB3 monochrome camera (Blackfly® BFLY-U3-23S6M-C; Teledyne FLIR Integrated Imaging Solutions, Richmond, BC, Canada).

### Monster

In monster sessions, a plastic mask (18cm height, 18cm width, 5cm depth) was placed on a metal arm attached to a gear rack so that it protruded into the far end of the arena, facing the shelter. When a mouse reached 30cm into the arena (towards the monster and the water spout), a beam break triggered the monster to move forward or charge (10 cm at 20cm/s), during which time a loud complex tone (120 dB) played. Upon charging, the monster paused for 500ms, then retreated again at 20cm/s. The monster continued moving back and forth, with accompanying tone, until the mouse returned to the shelter and the door closed, at which point the monster and tone stopped at once. No-monster sessions consisted of the same trial structure and were conducted in the absence of the monster and crossing the 30 cm line did not result in the triggering of the loud complex tone.

### Training

Mice were singly housed following surgery in both the photometry and ablation cohorts. Mice were first habituated to the experimenter: the experimenter handled each mouse for 20 minutes on two subsequent days. On the second day, the mouse was placed on water restriction, which continued until behavioral sessions ended. On the third day, each mouse was placed in the shelter for 30 minutes with water and food on the floor, and the door to the arena closed. Next, the mouse entered the training period, which lasted until mice either reached expert performance (80% of trials rewarded) or a preset maximum number of training sessions (ablation experiments: 5 sessions; recording experiments: 3 sessions). During the training period mice completed one session/day, consisting of 10 (ablation cohort, 23 animals; photometry cohort 1, 5 animals) or 30 (photometry cohort 2, 9 animals) trials. In each session, the mouse started in the shelter with the door closed. Each trial started with the door opening automatically, and the mouse was free to enter. Entrance into the arena was defined as the time when a beam 5cm outside of the door was broken, and subsequent movements were logged with additional beam breaks. If the mouse found and licked the water spout, a drop of water (10μl) was released from the spout. While mice were free to continue licking the spout, they could only receive one drop of water per trial. A trial ended when the mouse returned to the shelter and the door closed automatically behind (triggered by a beam break inside the shelter). A 20-second intertrial interval then occurred until the door opened again for another trial. After the training sessions, mice received water every day from experimenters and their body weights were checked daily to ensure that they were above 85% of their weight with freely available water (as measured one day before start of water restriction).

### Behavioral tests

Once mice had completed training sessions with a success rate of at least 80%, they progressed to the Monster Task sequence, in which no-monster sessions (no monster present, identical to training sessions) alternated with monster sessions (monster present). Mice in the ablation cohort moved through three rounds of no-monster and monster sessions, for a total of six days of testing. Mice in the photometry cohorts moved through one round of no-monster and monster sessions, for a total of two days of testing.

### Behavioral analysis

In the monster session, two types of failure were possible: failure before the monster was activated (e.g., before the mouse crossed the 30cm threshold) and failure after the monster was activated (after the monster crossed the 30cm threshold. For each session, success rate was calculated as the fraction of trials on which the animal succeeded. To average across mice, each mouse’s individual success rate was first calculated and then the average was taken across mice. On monster trials, we additionally calculated two distinct failure rates: first, the rate of failure before monster activation (e.g., before the mouse crossed the 30cm line) and the rate of failure after monster activation (after the mouse crossed the 30cm line). The latter was calculated as the number of trials on which the monster was activated and the mouse failed, over the total number of trials on which the monster was activated, to control for intra-animal differences in the rate of monster activation.

### Surgical procedures

Surgeries were performed under aseptic conditions in accordance with Harvard FAS IACUC protocol. Animals were anesthetized with isoflurane with an initial induction at 4% (1.0 l/min) which was lowered to 1-2% once the animal’s respiratory rate stabilized. The following AP/ML/DV coordinates were used for targeting injections and/or implants: TS −1.7/3.0/-2.3, VS 1.2/1.4/4.2^74^.

### Bilateral ablation of dopamine neurons

Bilateral ablation of dopamine neurons projecting to TS or VS was performed according to an existing protocol utilizing 6-hydroxydopamine (6-OHDA), a neurotoxin that is taken up through dopamine transporters^75^ and progressively kills the recipient neurons^9,10,76^. We first injected a mixture of a norepinephrine reuptake inhibitor (desipramine) and a monoamine oxidase inhibitor (pargyline) to prevent uptake of 6-OHDA by noradrenergic neurons and to prolong the viability of the 6-OHDA, respectively. Specifically, 28.5 mg desipramine and 6.2 mg pargyline were dissolved in 10mL of 0.9% NaCL at 45 °C, and then neutralized to pH 7.4 with NaOH; this mixture was injected at 10ml/kg intraperitoneally 30 minutes before intracranial injection of 6- OHDA. For preparation of 6-OHDA, according to extant protocol we first prepared a solution of 0.2% (2mg/ml) ascorbic acid in 5mL of 0.9% NaCl made with degassed water. Then, 9.15mg 6- OHDA HCl was dissolved in 0.5mL of the 0.2% ascorbic acid solution. The solution was kept on ice, wrapped in aluminum foil. It was discarded and made fresh if the solution turned red or brown (indicating that the 6-OHDA had broken down), or after 3 hours. Mice were given at least one week, and up to two weeks, following surgery for recovery. During this time, ketoprofen and buprenorphine were administered as an analgesic, with supplementary provisions of saline and a heating pad if the animal showed signs of dehydration (as assessed by daily checks).

### Fluorometry (photometry) surgical procedure

We used a triple transgenic mice (DAT-Cre+/-; GCaMP6f+/-; tdTomato+/-) (n = 11) so all mice genetically expressed GCaMP6f under the DAT promoter (see above). In all mice used for photometry recordings, we implanted an optic fiber (200 μm diameter, NA .37, RM2_FLT, Doric Lenses, Canada) into the target regions, using the above-specified coordinates, with the fiber tilted at 10 degrees from the plane of the skull surface. The fiber was attached to a custom stereotactic adapter and lowered slowly into the target site. Attachment to the skull was first secured with cyanoacrylate adhesive. Once solidified, this adhesive was followed by an opaque dental cement (Metabond) mixed with charcoal powder. Once this final sealant was dry (15-30 minutes), the surgery was complete.

### Histology and immunohistochemistry

Following completion of testing, brains were collected after perfusion with saline (PBS) followed by 4% paraformaldehyde (PFA). After 24 hours or more, the brain was sliced into 100 µm thick coronal sections using a vibratome (Leica, Germany) and stored in PBS. Visualization of dopamine axons in the striatum was accomplished through immunohistochemistry.

Specifically, brain sections were first incubated with rabbit anti-tyrosine hydroxylase (TH) antibody (AB152, Millipore Sigma, MO) at 4°C overnight, and then with a fluorescent goat anti-rabbit secondary antibody (A-11012, Thermo Fisher Scientific, MA) at 4°C for 2-12 hours.

Slices were then mounted in 4’,6-diamidino-2-pehnylindole (DAPI)-containing anti-fade solution (VECTASHIELD anti-fade mounting medium, H-1200, Vector Laboratories, CA) and imaged with a Zeiss Axio Scan Z1 slide scanner fluorescence microscope (Zeiss, Germany) at the Harvard Center for Biological Imaging.

### Fluorometry (photometry) recording

Fiber photometry was performed as previously reported using a bundle-imaging photometry setup (BFMC6_LED(410-420)_LED(460-490)_CAM(500-550)_LED(555_570)_CAM(580-680)_FC, Doric Lenses)^77^. This system collects fluorescence emitted through a low autofluorescence branching bundle patchcord (BBP(3)_200/220/900-0.37_3m_FCM- 3xCM2_LAF). Imaging the top of the branched bundle allowed simultaneous recording of signals emitted from multiple implant sites. 473 nm excitation light and 560 nm excitation light (LEDs from Doric Lenses) were delivered in an interleaved fashion for independent collection of fluorescence from GCaMP6f and tdTomato. GCaMP-emitted fluorescence fluctuates with intracellular calcium in cells expressing DAT, while tdTomato-emitted fluorescence is independent of calcium changes and thus serves as a control for motion-induced or other artifactual fluctuations in the photometry signal which are unrelated to calcium flux. We acquired GCaMP and tdTomato-emitted signals at 12Hz or 20Hz. Light intensity was calibrated before each cohort of mice to an intensity of 100μW, then held constant throughout all recording sessions for that cohort (1-2 weeks). Recording sessions were separated by at least 48 hours to ensure recovery of the signal from photobleaching. The Doric system also received a synchronizing pulse from the behavioral task software which allowed for temporal alignment of the photometry and behavioral data.

### Signal analysis

GCaMP (green) and tdTomato (red) signals were collected via separate CMOS cameras which simultaneously captured activity (as reflected by greyscale pixel intensity) across all regions of interest (ROIs) imaged. Signals were passed through a low-pass filter which removed signals above 1Hz. Next, signals were smoothed with a 50ms window, and global signal decay was normalized with a moving median, using a 100s window. Next fitted red signals were subtracted from green signals if the two signals were significantly correlated. Finally, the corrected signal was Z-scored using activity across the full session to calculate mean and standard deviation. When examining responses aligned to a specific behavioral event, the average activity 2s before the event was subtracted to account for extant elevations in activity unrelated to the event. To prevent large phasic reward responses from confounding our kinematic analyses, we removed reward-evoked fluorescence transients using a kernel-regression approach: we modeled the reward response as a smooth, time-varying impulse-response kernel and regressed this predicted reward-locked component out of the raw fluorescence trace. For each animal, we constructed a set of evenly spaced raised-cosine basis functions spanning a fixed peri-event window around reward delivery (−2 to +2 s). Basis functions were multiplied by a parabolic taper to down-weight the window edges and then L2-normalized to place all bases on a comparable scale. Reward delivery times were encoded as an impulse train on the same sampling grid as the photometry trace and each basis function was convolved with this impulse train to generate a set of reward regressors. We then fit a linear model to the instantaneous z-scored dopamine trace using the reward regressors to estimate the reward kernel coefficients. The fitted reward component at each time point was computed from the design matrix and estimated coefficients, and the reward-removed dopamine trace was defined as the residual. Besides event-locked activity analyses, all other analyses used this reward-removed photometry trace so that kinematic coupling (and other continuous relationships) could be assessed without contamination from reward delivery-evoked responses.

### DeepLabCut tracking

We used DeepLabCut version 2^78^ to track the mouse’s movement frame-by-frame throughout the arena. We tracked 7 mouse body parts (nose, left ear, right ear, upper back, lower back, and tail stem). After analyzing each behavioral session’s video with the DeepLabCut network, we used the generated output files (csv array containing x and y coordinates and likelihood values for each body part) to find and correct aberrant labels. To correct transient mistracks, we applied an automated cleaning procedure separately for each body part that flagged frames as invalid if tracking confidence fell below a fixed likelihood threshold (0.85) or if the frame-to-frame displacement exceeded an adaptive “jump” threshold. The jump threshold was estimated from high-confidence frames only and set to the mean displacement plus 1.5 standard deviations, allowing the cutoff to scale with each animal/body part’s typical movement. Invalid samples were replaced with missing values and then linearly interpolated across time (with extrapolation at the beginning/end if needed) to produce continuous x/y traces. To reduce noise from any single marker, the animal’s planar position was defined as the average of three head markers (nose, left ear, right ear). The head base was defined as the midpoint between the two ears.

Because video (84 Hz) and photometry (12 Hz or 20 Hz) were sampled at different rates, the video-derived position and marker traces were resampled within each trial to match the number of photometry samples in the corresponding interval, enabling sample-wise alignment between behavior and neural activity.

#### Velocity, acceleration, and scalar kinematics

Cartesian velocity and acceleration were computed by finite differences.

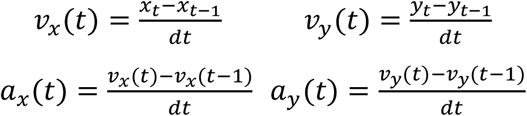

Speed and scalar acceleration were defined as the Euclidean magnitude of the instantaneous velocities and accelerations, respectively.

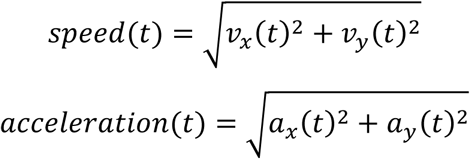

To decompose motion with respect to the water spout, we defined the position vector relative to the spout.

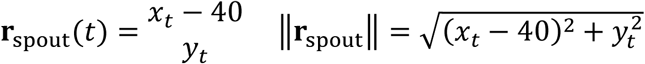

Radial velocity was then computed as the projection of 𝑣 onto **r**.

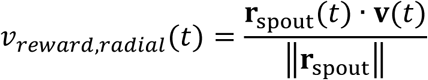

### Statistical analysis

Experimenters were blinded to the condition of animals in the dopamine ablation studies until behavioral experiments were complete. Power analysis was used to determine the sample size (n=12) necessary to detect a significant difference in success rate during control (reward-only) foraging based on an initial sample of 6 controls and 8 animals with bilateral ablations of dopamine neurons projecting to the ventral striatum.

For comparing average continuous measures between groups, we used t-tests. To analyze repeated-measures data, we used mixed-effects models that accounted for within-animal dependence (and, where applicable, multiple recording sites per animal). Binary outcomes (e.g., success rate, avoidance) were analyzed with generalized linear mixed-effects models using a logit link, whereas continuous outcomes were analyzed with linear mixed-effects models. For photometry analyses, mixed-effects models were used to quantify relationships between dopamine activity and behavioral variables. Post hoc contrasts, estimated marginal means, and estimated marginal trends were computed using emmeans, and *p*-values were adjusted for multiple comparisons where appropriate.

### Survival Analyses

We quantified persistence to approach the reward using survival analysis (Figure 1G). For each subject, we defined the distance variable as the maximum approach distance reached on a trial (bounded between 0–40 cm) and an event indicator marking whether the animal terminated the approach before reaching the water spout. We then constructed a Kaplan–Meier estimator of the probability of continuing approach as a function of distance and fit separate survival curves by ablation group using survfit from the R survival package. Group differences were evaluated with a log-rank test, and results were plotted using survminer.

To quantify group differences in the probability to abandon reward approach as animals progressed toward the spout, we fit Cox proportional hazards models with distance as the analysis axis and quitting as the event. Trials in which animals did not quit before the spout were treated as right-censored observations. Ablation status was included as a predictor, and to allow the ablation effect to vary over distance we incorporated a time-varying ablation term parameterized as an interaction with a log-transformed distance function. Baseline hazards were allowed to differ across sessions using stratification, and repeated observations within animals were accounted for using an animal-level frailty term. From the fitted model, we computed a distance-dependent hazard ratio comparing ablation to control across the spout approach range by evaluating the ablation effect at each distance and exponentiating the resulting log-hazard difference. Confidence intervals for the hazard ratio as a function of distance were obtained by propagating uncertainty from the model’s variance-covariance matrix to the distance-dependent log-hazard ratio and transforming the ±1.96 standard error bounds. Hazard ratios greater than 1 indicate increased instantaneous risk of quitting in the ablation group relative to controls, whereas values less than 1 indicate reduced quitting risk.

### Event-locked significance testing

To assess whether dopamine activity showed a significant event-locked response, we used a circular-shift permutation test within each recording session (Figure 3D). For each animal, session, hemisphere, region (VS, TS), and outcome (rewarded, unrewarded), we identified all occurrences of each event type (Door Open, Arena Entrance, Monster Charge, Reward Delivery, Shelter Entrance). For each event occurrence, we quantified the response magnitude as the peak Z-scored fluorescence in a fixed post-event window (0–1 s after event onset). The observed event-locked statistic for a given event type was defined as the mean of these per-event post-event peaks within that subset. To generate a null distribution that preserved the autocorrelation and amplitude statistics of the photometry trace while disrupting alignment to event times, we circularly shifted the fluorescence signal by a random offset within the session and recomputed the same mean post-event peak statistic. Shift offsets were constrained to be at least 2.1 s (i.e., longer than the peri-event window) to avoid trivial near-alignments. This procedure was repeated 2,000 times per subset and event type to generate a null distribution of mean post-event peak responses expected by chance under preserved signal autocorrelation. We summarized the event-locked response magnitude as a null-corrected statistic defined as the observed mean post-event peak minus the median of the null distribution. Permutation-derived null-corrected effect sizes were analyzed at the population level using linear mixed-effects models to test how event-locked response magnitude depended on event, outcome, and region. Estimated marginal means were computed using emmeans, and contrasts were Bonferroni-corrected for multiple comparisons.

### Kinetic and reward models

We quantified how well kinematic variables explained dopamine fluorescence using linear mixed-effects models (Figure 4C). Prior to fitting, Animal, Session, Hemisphere, Region (VS, TS), and Trial were treated as categorical factors. Dopamine activity (𝑍) was modeled with region-specific fixed effects by interacting each predictor with Region, yielding separate slopes for each region. To account for repeated measurements, we included random intercepts for Animal and for recording site nested within animal (Animal:Site). We reduced temporal autocorrelation in two steps. First, we downsampled the photometry and kinematic signals to 1 Hz by averaging within 1 s bins to attenuate fast-timescale correlations and yield observations at the approximately at the scale of our behavioral regressors. Second, we included a one-step lag of the dopamine signal (𝑍_𝑡−1_) in the models to capture remaining first-order temporal persistence, verifying adequacy by confirming that residual autocorrelation was minimal beyond lag 0. This approach was used for all model-based analyses relating photometry signals to kinematics.

We first fit a “movement-only” model containing z-scored speed, acceleration, and angular velocity:

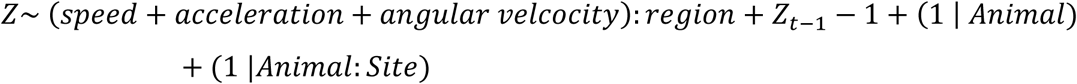

We then fit an expanded model that additionally included z-scored distance to spout and reward-referenced kinematic variables (radial velocity toward the spout, radial acceleration, and angular velocity in the reward-referenced frame):

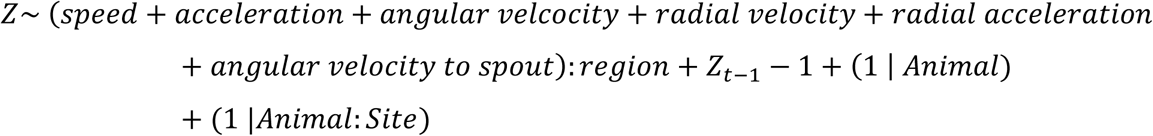

Both models were fit using maximum likelihood to enable nested model comparison. We compared the expanded and movement-only models using a likelihood ratio test (ANOVA) and information criteria (AIC and BIC). Overall model fit statistics were additionally summarized using the performance package.

### Defining Approach vs Retreat

To derive approach and retreat movement epochs, X and Y positions were smoothed using a Savitzky–Golay filter with an adaptive window based on sampling rate (∼1.5 s). The derivative of the smoothed X-position was estimated via a Savitzky–Golay derivative and then low-pass filtered (Butterworth; default cutoff 0.3 Hz, applied with zero-phase filtering). Behavioral state was labeled from smoothed position along the shelter to spout axis (X) and its derivative: samples were categorized into distal and proximal approach sub-bands based on X-position thresholds, with Retreat overriding these labels when the smoothed X-derivative indicated movement back toward shelter. Periods around reward consumption and exploration beyond the spout were labeled separately and excluded from analyses.

### Value Maps

To compute the neural “value” signal used for mapping (Figure 6), dopamine activity (Z) was converted to a within-epoch cumulative integral using a Riemann rule:

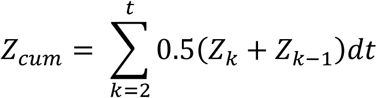

The resulting cumulative trace was z-scored within the same grouping. Value maps were then computed by binning position along the shelter–spout axis and averaging 𝑍_𝑐𝑢𝑚_ within bins at the animal level and trial level. To generate a null control, the signal was circularly permuted within trial for 1000 permutations and the same bin-averaged profiles were recomputed to produce shuffle-derived distributions and mean shuffled maps. To quantify map structure, monotonicity metrics were extracted from the binned profiles, including Spearman correlation between binned X and mean signal per animal, and (the number/fraction of increasing steps across adjacent bins above a small slope threshold (0.02).

## Data availability

All data (both behavioral and fluorometry) will be deposited in Dryad. All analysis codes will be available at Github.

## Acknowledgements

We thank Melissa Tian for histology summary and all lab members for discussion. We thank the Harvard Center for Biological Imaging for infrastructure and support. This work was supported by grants from National Institute of Health (F30MH132298, I.G.; T32GM007753, I.G.; U19NS113201, N.U.; R01MH133950, N.U., M.W.-U.; R01DA059751, N.U., M.W.-U.; R01MH125162, M.W.-U.), Simons Collaboration on the Global Brain (N.U.), and Bipolar Disorder Seed Grant Program (N.U.).

## Author Contributions

I.G., E.S.I. and M.W.-U. designed the experiments and analyzed the data. I.G., E.S.I. and A.K. collected the data. The results were discussed and interpreted by I.G., E.S.I., N.U. and M.W.-U. I.G., E.S.I. and M.W.-U. wrote the paper and I.G., E.S.I., N.U. and M.W.-U. edited the paper.

## Declaration of Interests

The authors declare no competing interests.

---

**Figure S1.**
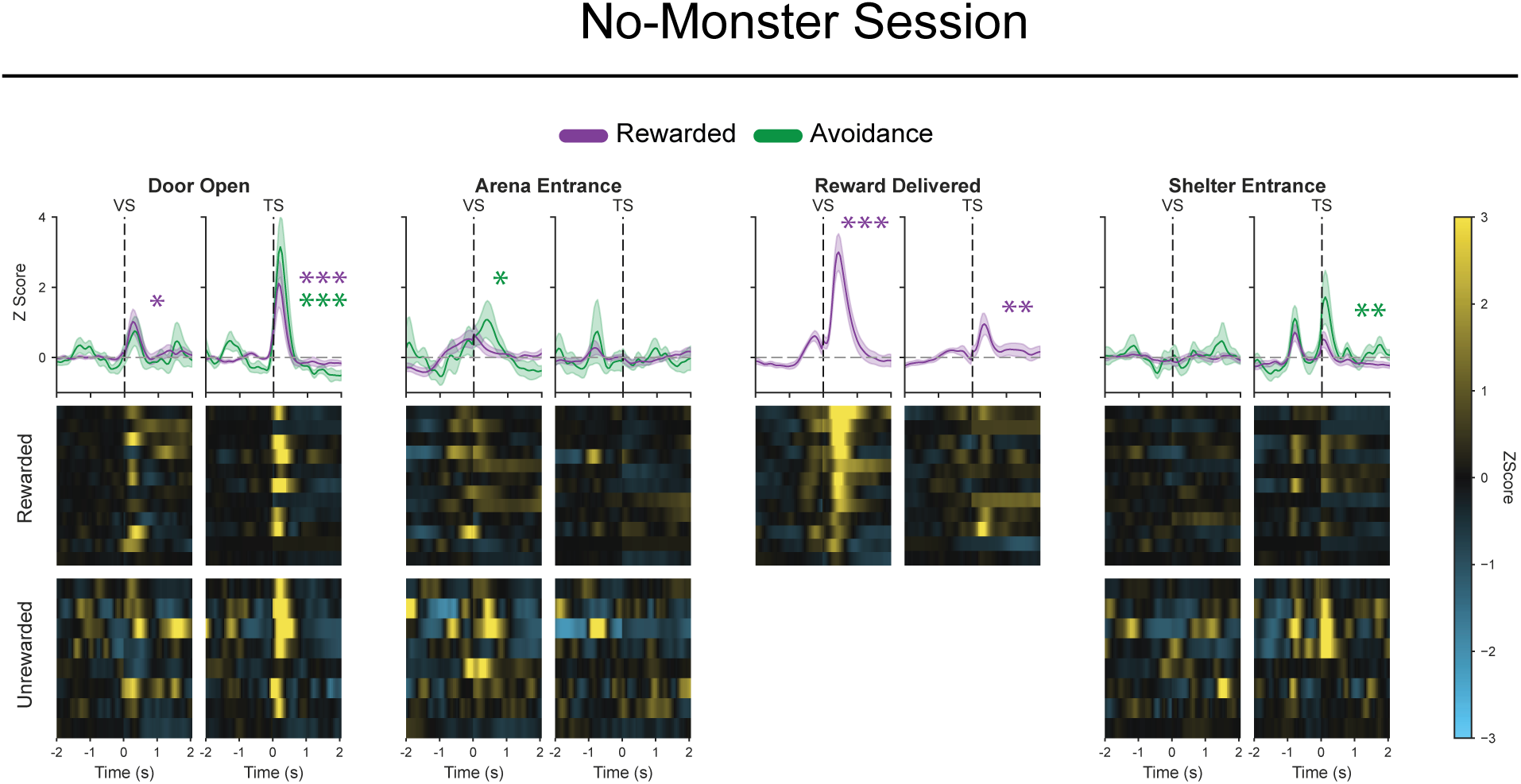
Fiber photometry recordings from ventral and tail of striatum dopamine axons in no-monster sessions. Related to. Figure 3. Event-aligned dopamine axon signals (z-scored) in VS and TS around key task events in reward acquisition (purple) and avoidance (green) trials: door opening (one-sample t-test: TS, reward, *t*(99.3)=5.7, *p* = 2.1 × 10^-7^; avoidance, *t*(128.0)=7.1, *p* = 1.6 × 10^-10^; pairwise t-test: reward–avoidance *t*(185)=−1.7, *p*=0.075; one-sample t-test: VS, reward, *t*(152.5)=2.7, *p*=0.015; avoidance, *t*(181.6)=1.4, *p*=0.31; pairwise t-test: reward– avoidance, *t*(186)=0.75, *p*=0.45), arena entrance (one-sample t-test: TS, reward, *t*(99.3)=−0.54, *p*=1.00; avoidance, *t*(128.0)=0.28, *p*=1.00; pairwise t-test: reward–avoidance *t*(185)=−0.66, *p*=0.51; one-sample t-test: VS, reward, *t*(152.5)=1.52, *p*=0.26; avoidance, *t*(181.6)=2.5, *p*=0.021; pairwise t-test: reward–avoidance *t*(186)=−1.0, *p*=0.28), reward delivery (one-sample t-test: TS, reward, *t*(99.3)=2.65, *p*=0.0093; VS, reward, *t*(152.5)=7.1, *p* = 4.4 × 10^-11^), and shelter entrance (one-sample t-test: TS, reward, *t*(99.3)=0.83, *p*=0.82; avoidance, *t*(128.0)=3.3, *p*=0.0022; pairwise t-test: reward–avoidance, *t*(185)=−2.2, *p*=0.024. one-sample t-test: VS, reward, *t*(152.5)=−0.13, *p*=1.00; avoidance, *t*(181.6)=0.30, *p*=1.00; pairwise t-test: reward–avoidance *t*(186)=−0.35, *p*=0.72). Mean ± SEM. Colored asterisks denote significant deviations from baseline. n = 12 animals. * *p* < 0.05, ** *p* < 0.01, *** *p* < 0.001.

**Figure S2.**
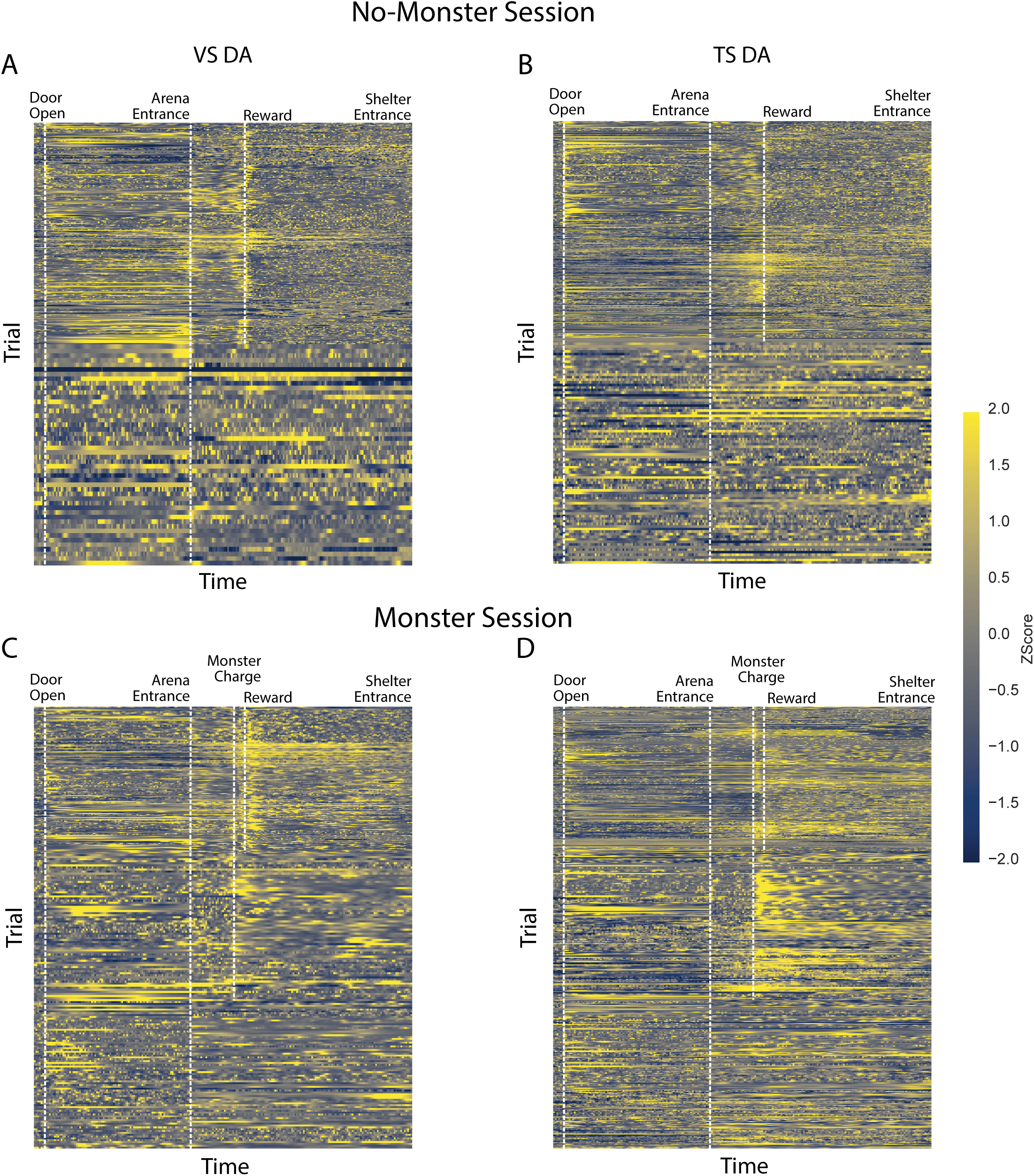
Dopamine axon activity stretched and aligned to key behavioral events in no- monster and monster sessions. Related to. Figure 3. (A–B) Z-scored dopamine axon activity in VS (A, C) and TS (B, D) in all trials (rows) in no-monster sessions (A-B) and monster sessions (C-D) in all animals. To enable visual comparison across trials with variable durations of events, dopamine axon activity in each trial was resampled to a common normalized trial axis spanning door opening through shelter entrance, preserving the order of within-trial events. Vertical dashed lines mark the mean-aligned positions of key events.

**Figure S3.**
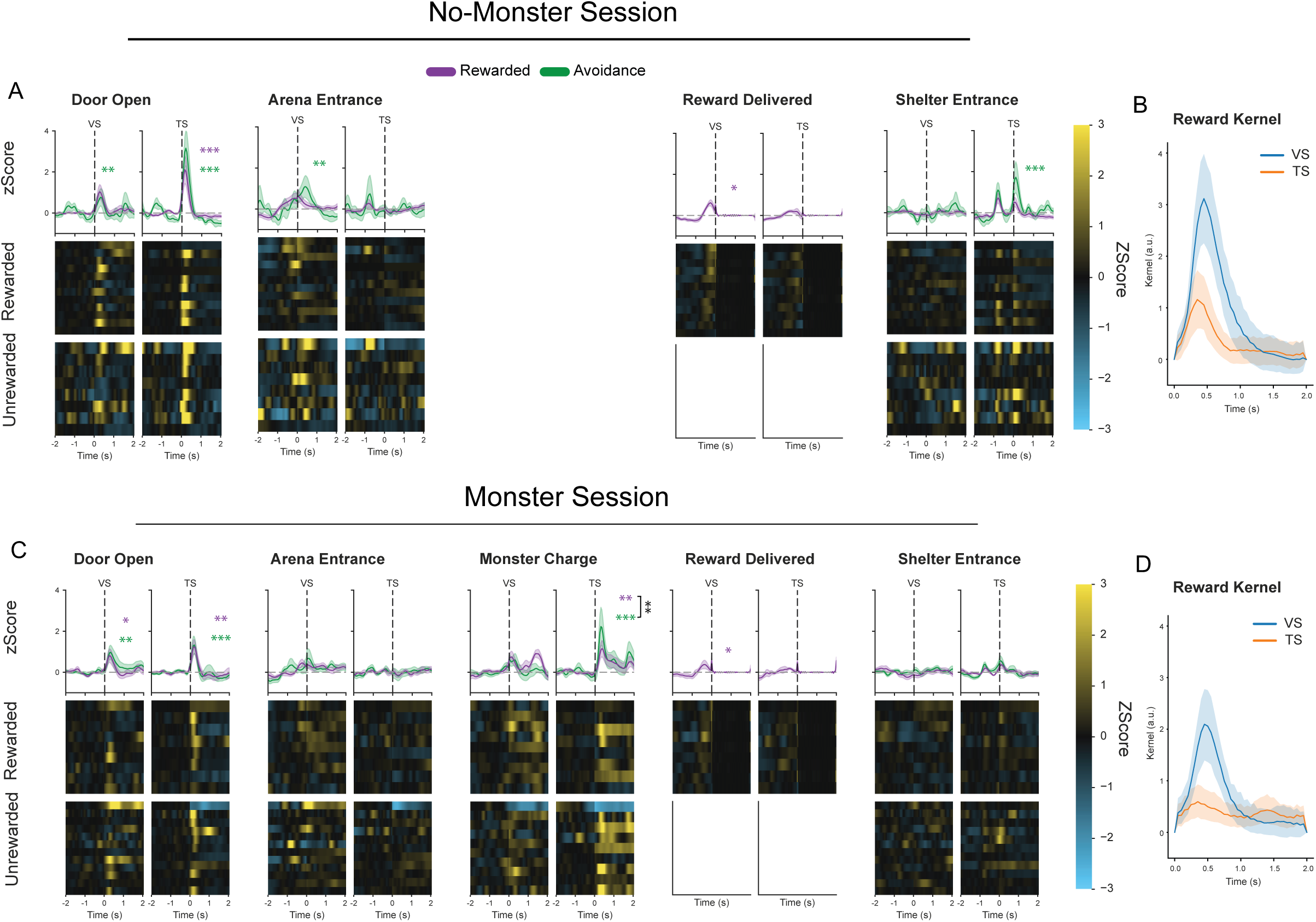
Validation of kernel-regression for phasic reward response removal in VS and TS dopamine axon recordings. Related to. Figure 3. Similar plots as Figure 3 but removing responses at reward delivery (see Methods). (A) Event-aligned signals (z-scored) for reward acquisition (purple, “reward”) and avoidance (green, “avoidance”) trials in dopamine axons in VS and TS after removing phasic responses to reward delivery in the no-monster session at door opening (one-sample t-test: TS, reward, *t*(95.80) = 6.35, *p* = 1.43 × 10^-8^; avoidance, *t*(124.39) = 7.73, *p* = 6.32 × 10^-12^; VS, reward, *t*(148.16) = 2.97, *p* = 0.0070; avoidance, *t*(178.52) = 1.61, *p* = 0.22; pairwise t-test: TS, reward vs avoidance, *t*(184.85) = −1.87, *p* = 0.063; VS, reward vs avoidance, *t*(185.96) = 0.81, *p* = 0.42), arena entrance (one-sample t-test: TS, reward, *t*(95.80) = −0.52, *p* = 1.00; avoidance, *t*(124.39) = 0.26, *p* = 1.00; VS, reward, *t*(148.16) = 1.68, *p* = 0.19; avoidance, *t*(178.52) = 2.86, *p* = 0.0095; pairwise t-test: TS, reward vs avoidance, *t*(184.85) = −0.63, *p* = 0.53; VS, reward vs avoidance, *t*(185.96) = −1.20, *p* = 0.23), reward delivery (one-sample t-test: TS, *t*(95.80) = 1.10, *p* = 0.27; VS, *t*(148.16) = 2.61, *p* = 0.010), shelter entrance (one-sample t-test: TS, reward, *t*(95.80) = 0.94, *p* = 0.70; avoidance, *t*(124.39) = 3.61, *p* = 0.00087; VS, reward, *t*(148.16) = −0.15, *p* = 1.00; avoidance, *t*(178.52) = 0.37, *p* = 1.00; pairwise t-test: TS, reward vs avoidance, *t*(184.85) = −2.44, *p* = 0.016; VS, reward vs avoidance, *t*(185.96) = −0.43, *p* = 0.67). (B) Reward kernel for VS (blue) and TS (orange) in the no-monster session. (C) Event-aligned signals (z-scored) for reward acquisition (purple, “reward”) and avoidance (green, “avoidance”) trials in dopamine axons in VS and TS after removing phasic responses to reward delivery in the monster session at door opening (one-sample t-test: TS, reward, *t*(193.75) = 2.97, *p* = 0.0066; avoidance, *t*(160.89) = 4.12, *p* = 0.00012; VS, reward, *t*(237.00) = 2.32, *p* = 0.042; avoidance, *t*(213.09) = 3.43, *p* = 0.0015; pairwise t-test: TS, reward vs avoidance, *t*(248.45) = −0.55, *p* = 0.58; VS, reward vs avoidance, *t*(247.94) = −0.62, *p* = 0.54), arena entrance (one-sample t-test: TS, reward, *t*(193.75) = −0.26, *p* = 1.00; avoidance, *t*(160.89) = −0.37, *p* = 1.00; VS, reward, *t*(237.00) = 1.03, *p* = 0.60; avoidance, *t*(213.09) = 1.58, *p* = 0.23; pairwise t-test: TS, reward vs avoidance, *t*(248.45) = 0.055, *p* = 0.96; VS, reward vs avoidance, *t*(247.94) = −0.31, *p* = 0.76), monster charge (one-sample t-test: TS, reward, *t*(193.75) = 3.43, *p* = 0.0015; avoidance, *t*(196.85) = 7.68, *p* = 1.43 × 10^-12^; VS, reward, *t*(237.00) = 1.84, *p* = 0.13; avoidance, *t*(240.33) = 1.52, *p* = 0.26; pairwise t-test: TS, reward vs avoidance, *t*(253.19) = −3.26, *p* = 0.0013; VS, reward vs avoidance, *t*(252.29) = 0.26, *p* = 0.80), reward delivery (one-sample t-test: TS, *t*(193.75) = 1.59, *p* = 0.11; VS, *t*(237.00) = 2.32, *p* = 0.021), shelter entrance (one-sample t-test: TS, reward, *t*(193.75) = 0.52, *p* = 1.00; avoidance, *t*(160.89) = 1.32, *p* = 0.38; VS, reward, *t*(237.00) = −0.45, *p* = 1.00; avoidance, *t*(213.09) = 0.20, *p* = 1.00; pairwise t-test: TS, reward vs avoidance, *t*(248.45) = −0.53, *p* = 0.59; VS, reward vs avoidance, *t*(247.94) = −0.52, *p* = 0.60). (D) R ward kernel for VS (blue) and TS (orange) in the monster session. Mean ± SEM across animals. n = 12 animals. *p* < 0.05, ** *p* < 0.01, *** *p* < 0.001.

**Figure S4.**
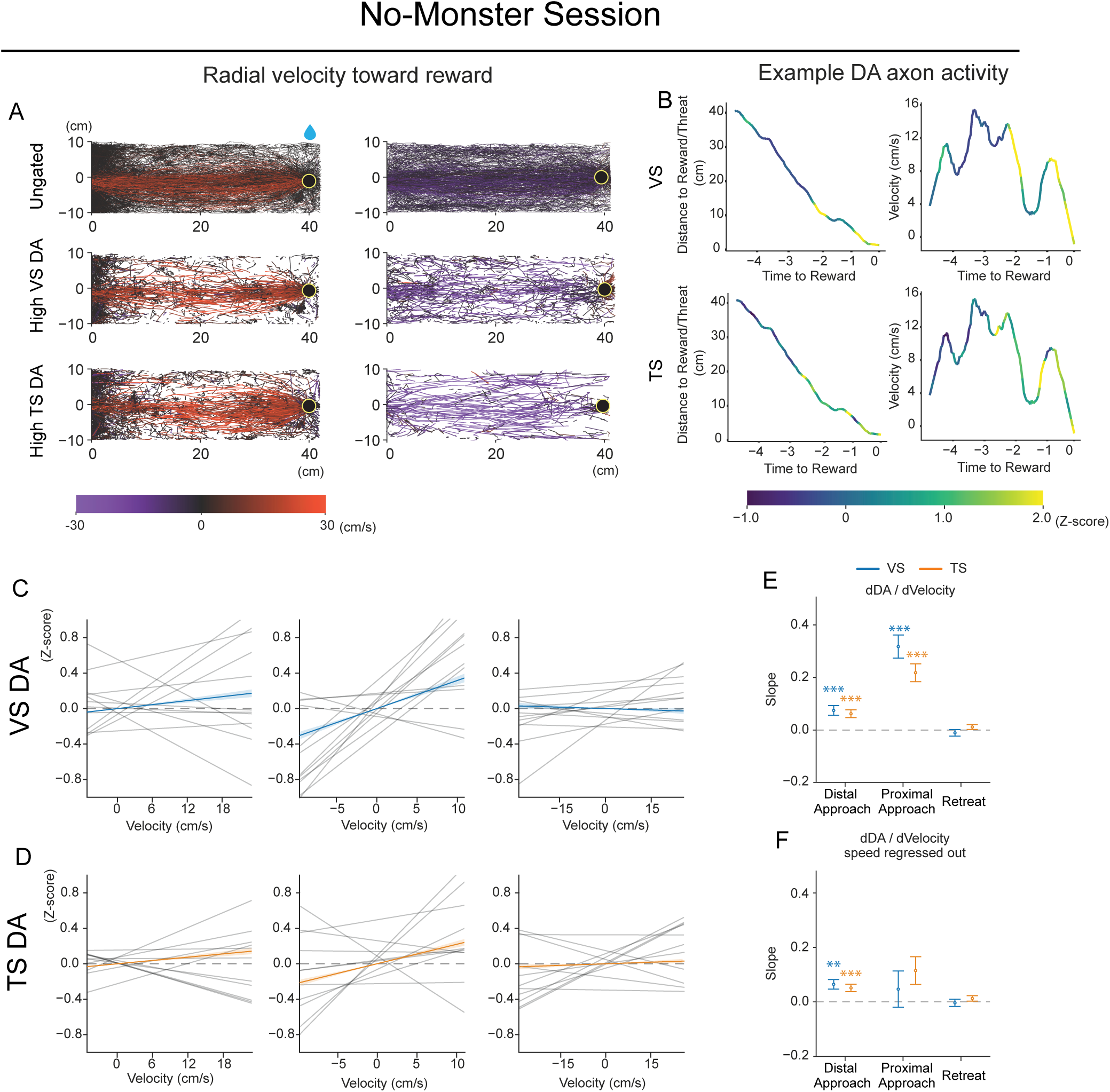
Correlation of VS and TS dopamine axon activity with reward approach kinematics in no-monster sessions. Related to. Figure 5. The same analyses as Figure 5 but for no-monster sessions. (A) Top, example animal trajectories during approach (left) and retreat (right) epochs, color-coded by radial velocity relative to reward/threat position (positive values indicate movement toward reward/threat; negative values indicate movement toward the shelter). Middle, trajectories from time points with high VS dopamine axon activity (Z > 1) during approach and retreat. Bottom, trajectories from time points with high TS dopamine axon activity (Z > 1) during approach and retreat. (B) Example of how VS (top) and TS (bottom) axonal activity varies with distance (left) and radial velocity (right) to reward in an exemplar mouse in a single trial. (C, D) Correlation between animal radial velocity toward reward and dopamine axon activity in VS (C) and TS (D). Thin black lines, per-animal regression slopes. Colored lines (blue, VS; orange, TS), the population-level mixed-effects regression slope ± SEM. (E) Estimated slopes for VS (blue) and TS (orange) dopamine axon activity vs radial velocity (dDA/ dVelocity) across behavioral epochs (one-sample Wald z-test: VS: distal approach, *z* = 4.04, *p* = 5.4 × 10^-4^; proximal approach, *z* = 7.2, *p* = 5.4 × 10^-12^; retreat, *z* = −0.86, *p* = 1.00; TS: distal approach, *z* = 4.22, *p* = 2.4 × 10^-4^; proximal approach, *z* = 6.43, *p* = 1.3 × 10^-9^; retreat, *z* = 1.25, *p* = 1.00) (F) Estimated slopes of dopamine axon activity versus radial velocity (dDA/dVelocity), with speed regressed out, across behavioral epochs (one-sample Wald z-test: VS: distal approach, *z* = 3.880, *p* = 0.0010; proximal approach, *z* = 0.76, *p* = 1.00; retreat, *z* = 0.012, *p* = 1.00; TS: distal approach, *z* = 4.00, *p* = 6.3 × 10^-4^; proximal approach, *z* = 2.34, *p* = 0.19; retreat, *z* = 1.62, *p* = 1.00). Open circles indicate the mean; error bars indicate ± SEM. n = 12 animals. * *p* < 0.05, *** *p* < 0.001.

**Figure S5.**
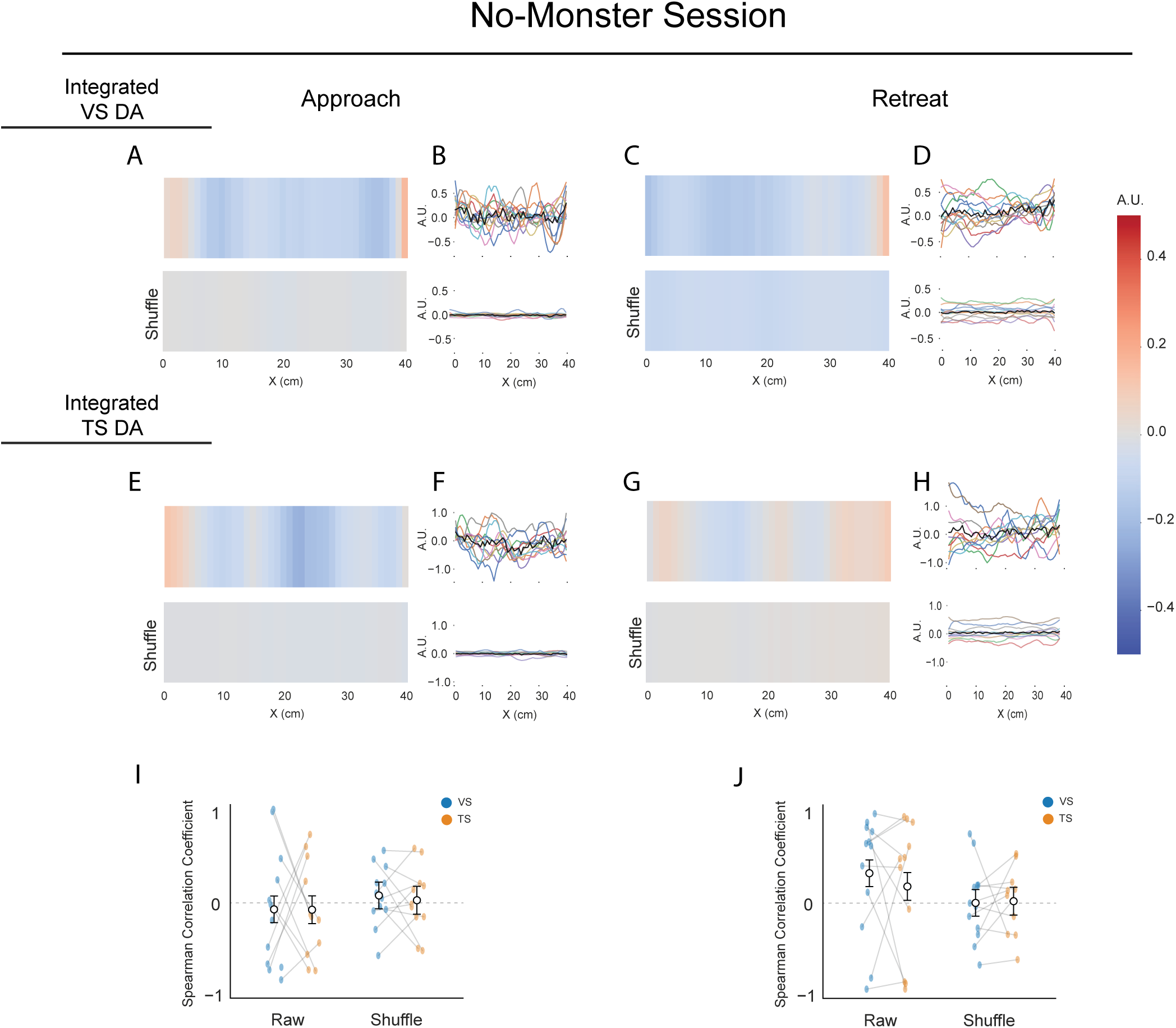
Integrated VS dopamine signals do not form clear spatial gradients in the absence of threat. Related to Figure 6. The same analyses as Figure 7A-H but for no-monster sessions. Integrated dopamine signals in VS (A-D) and TS (E-H) plotted as a function of position in the arena during approach (A-B, E-F) and retreat (C-D, G-H). Heatmaps show the mean position-binned integrated signal for the observed data (top) and a within-trial circularly permuted control (bottom). Line plots show integrated traces across the arena in each animal (colored lines) with the group mean overlaid (black). (I) Monotonicity of the gradient during approach was tested with Spearman correlation. No mean Spearman correlation differed significantly from zero (pairwise t-test: VS, raw, *t*(168)=-0.82, *p*=1.00; shuffle, *t*(168)=0.23, *p*=1.00; TS, raw, *t*(168)=-0.81, *p*=1.00; shuffle *t*(168)=−0.12, *p*=1.00). (J) Monotonicity of the gradient during retreat was tested with Spearman correlation. No mean Spearman correlation differed significantly from zero (pairwise t-test: VS, raw, *t*(168)=2.23, *p*=0.11; shuffle, *t*(168)=0.0037, *p*=1.00; TS, raw, *t*(168)=1.18, *p*=0.95; shuffle *t*(168)=0.13, *p*=1.00). n = 12 animals.

**Figure S6.**
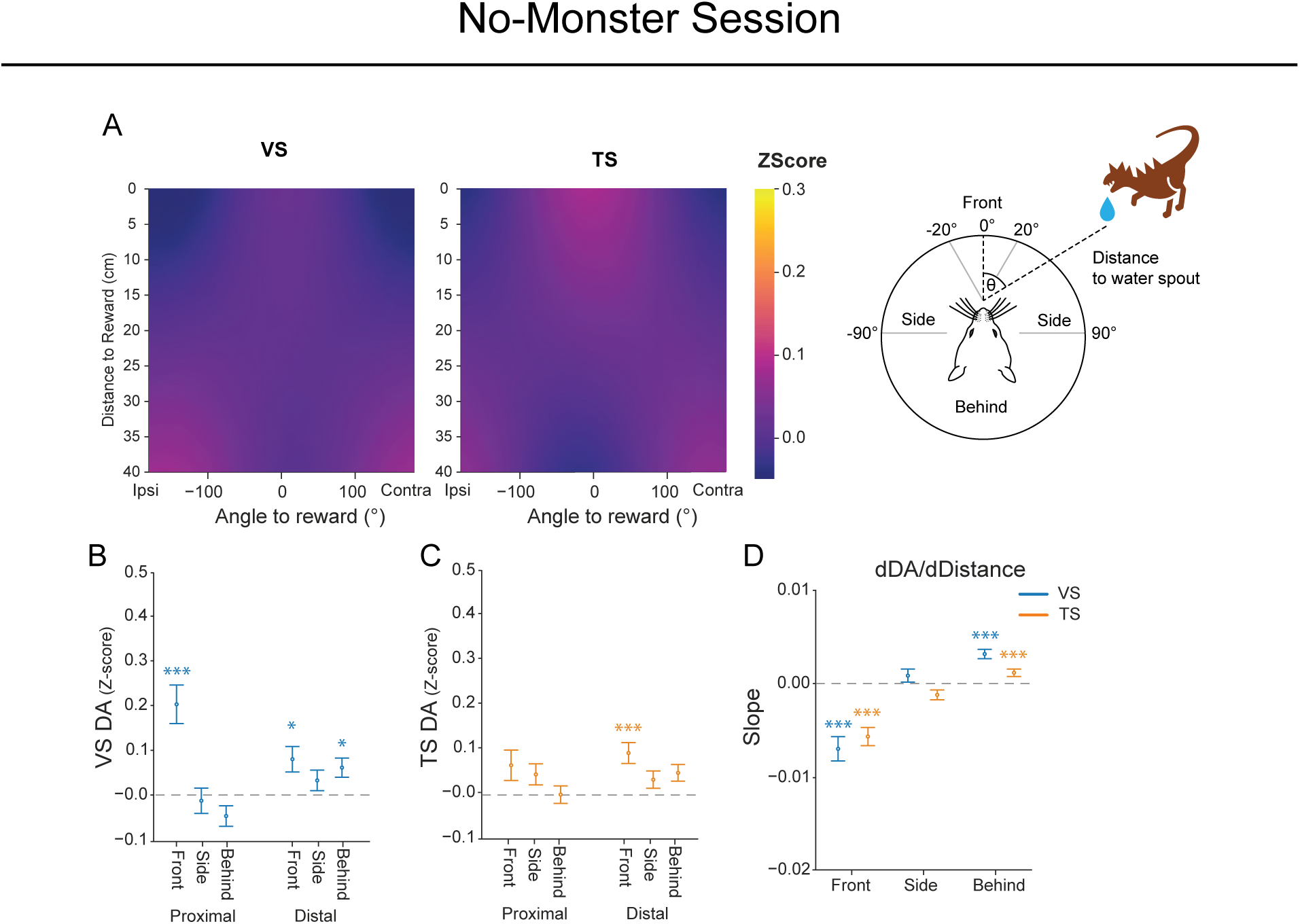
Correlation of VS and TS dopamine axon activity with reward proximity and direction in no-monster sessions. Related to. Figure 7. The same analyses as Figure 6 but for non-monster sessions. (A) Two-dimensional maps of z-scored dopamine axon activity in VS (left; pairwise t-test ipsi vs contra: *t*(822) = 0.53, *p*=0.60) and TS (right; pairwise t-test ipsi vs contra: *t*(143) = 0.86, *p*=0.39) as a function of head angle relative to and distance to reward. Color indicates model-estimated mean z-scored signals within each angle-distance bin. (B) Mean VS dopamine axon activity as a function of proximity (proximal vs distal) and orientation relative to the water spout (front, side, behind) (Wald z-test: proximal front, *z* = 4.8, *p* = 9.4 × 10^-6^; proximal side, *z* = −0.31, *p* = 1.0; proximal behind, *z* = −1.9, *p* = 0.36; distal front, *z* = 2.9, *p* = 0.019; distal side, *z* = 1.6, *p* = 0.66; distal behind, *z* = 3.0, *p* = 0.014). (C) Mean TS dopamine axon activity as a function of proximity and orientation to the reward/threat (Wald z-test: proximal front, *z* = 2.1, *p* = 0.23; proximal side, *z* = 2.1, *p* = 0.19; proximal behind, *z* = 0.24, *p* = 1.0; distal front, *z* = 4.2, *p* = 1.5 × 10^-4^; distal side, *z* = 1.9, *p* = 0.28; distal behind, *z* = 2.9, *p* = 0.024). (D) Slope of dopamine axon activity in VS (blue) and TS (orange) with respect to distance to reward (dZ/dDistance) within each orientation category (front, side, behind). Negative slopes indicate higher dopamine at positions closer to reward. (Wald z-test: VS: front, *z* = −5.3, *p* = 4.3 × 10^-7^; side, *z* = 1.2, *p* = 1.0; behind, *z* = 6.2, *p* = 2.8 × 10^-9^; TS: front, *z* = −5.9, *p* = 2.7 × 10^-8^; side, *z* = −2.3, *p* = 0.14; behind, *z* = 2.9, *p* = 0.022). Error bars, mean ± SEM. n = 12 animals. * *p* < 0.05, ** *p* < 0.01, *** *p* < 0.001.

**Figure S7.**
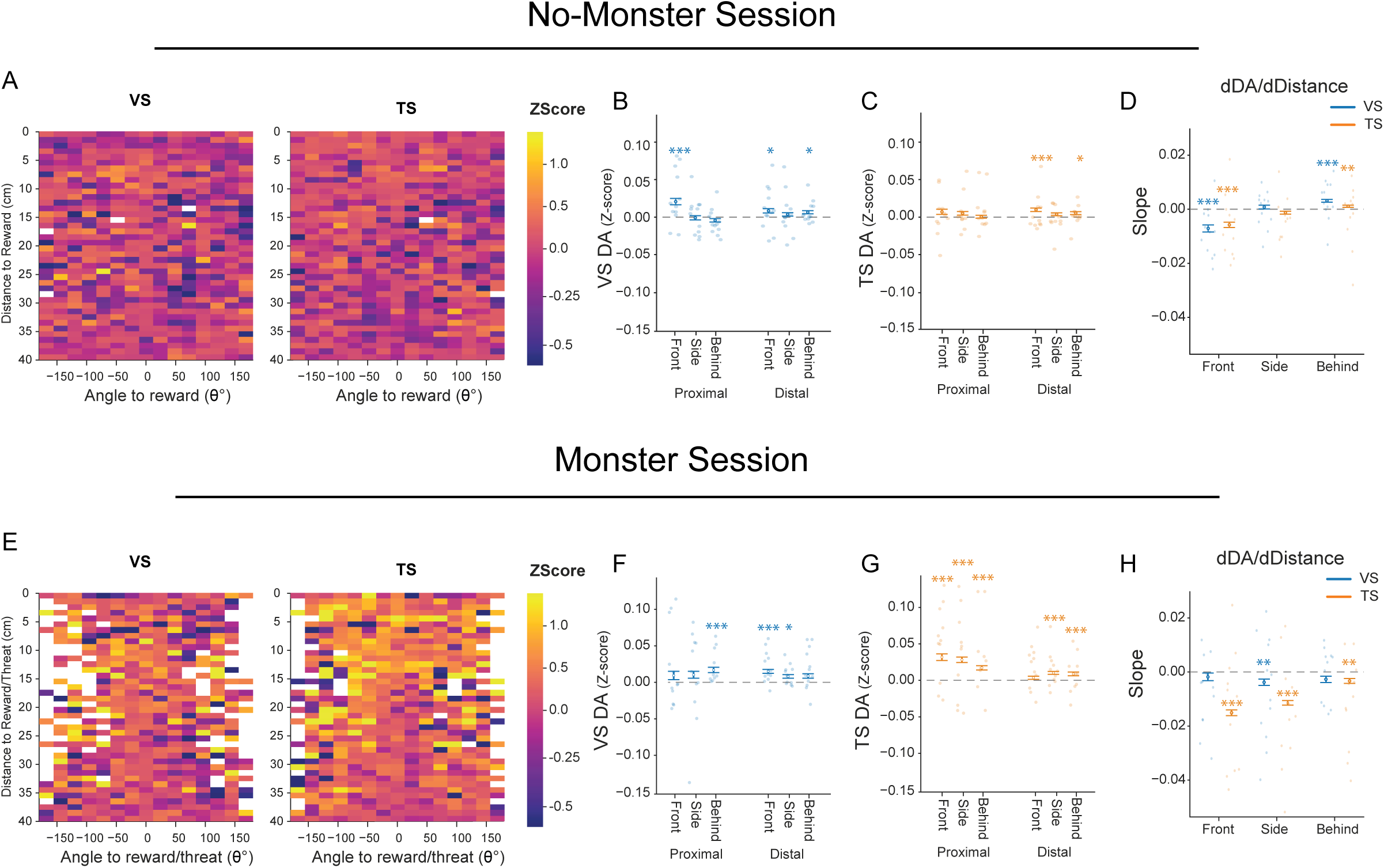
Animal level distribution of VS and TS dopamine axon activity modulation by proximity and direction to reward/threat. Related to. Figure 7. (A, E) The similar analyses as Figure 6A, but showing mean of original signals without model fitting in no-monster sessions (A) nd in monster sessions (E). Two-dimensional maps of z-scored dopamine axon activity in VS (Left) and TS (Right) as a function of head angle relative to and distance to reward. Color indicates mean z-scored signals within each angle-distance bin across animals. (B-D) The same plots as Figure S5B-D but with datapoints from individual animals (colored dots). (F-H) The same plots as Figure 6B-D but with datapoints from individual animals (colored dots). Error bars, mean ± SEM. n = 12 animals. * *p* < 0.05, ** *p* < 0.01, *** *p* < 0.001

## Notes

### Competing Interest Statement

The authors have declared no competing interest.

